# Transcriptomic and pathological analysis of the hnRNP network reveals glial involvement in FTLD pathological subtypes

**DOI:** 10.1101/2025.07.14.664732

**Authors:** A. Gatt, Y. Buhidma, K. Fodder, J. Humphrey, S.C. Foti, B. Frias, B.C. Benson, P. Gami-Patel, L.M. Gittings, C.E. Toomey, T. Lashley

## Abstract

Frontotemporal dementia (FTD) is a neurodegenerative disorder with a strong heritable component. Frontotemporal lobar degeneration (FTLD) refers to the pathological changes seen in FTD, characterised by atrophy of the frontal and temporal lobes and the presence of abnormal protein inclusions. In the case of FTLD with hyperphosphorylated TDP-43 positive inclusions (FTLD-TDP), five pathological subtypes (A, B, C, D, and E) are observed based on the types and distribution of inclusions found in the brain. In all subtypes, there tends to be a large variability in the number of pathological inclusions observed between cases, with limited correlation to clinical manifestations.

TDP-43 is an RNA binding protein belonging to the heterogeneous nuclear ribonucleoprotein (hnRNP) family which along with other hnRNPs modulates multiple aspects of RNA processing. HnRNPs other than TDP-43 have been implicated in several neurological diseases, including ALS, FTLD-TDP, FTLD-FUS and Alzheimer’s disease. Multiple hnRNPs have been found in pathological inclusions in specific subtypes of FTLD-TDP, suggesting potential roles in the disease process. The role of the hnRNP network in FTLD disease pathogenesis, however, has not yet been investigated. This study aimed to comprehensively evaluate the presence and expression of hnRNP proteins in two pathological subtypes of sporadic FTLD-TDP (A and C) as well as the genetic form FTLD-TDP A *C9orf72* using immunohistochemistry and gene expression analysis by single-nuclei RNA-sequencing.

We found that there was great variability in frequency of TDP-43 pathology across and within FTLD-TDP pathological subtypes. Finally, our findings suggest that distinct global transcriptomic profiles may underlie the different pathological subtypes of FTLD-TDP. The most prominent transcriptomic changes were observed in oligodendrocytes and astrocytes, involving multiple hnRNPs across FTLD subtypes compared to controls. Transcriptomic co-expression analysis further revealed that glial clusters were more strongly associated with RNA processing dysfunction and contribute to disease classification. Together, these findings highlight the involvement of the hnRNP network and glial-specific RNA processing alterations in FTLD-TDP pathophysiology, offering new insight into the molecular distinctions between pathological subtypes and potential targets for future investigation.

## Introduction

Frontotemporal dementia (FTD) is clinically categorised into three recognised syndromes known as behavioural variant frontotemporal dementia (bvFTD), semantic dementia (SD) and progressive non-fluent aphasia (PNFA). It can also overlap with motor neuron disease/ amyotrophic lateral sclerosis (MND/ ALS) (FTD-MND), corticobasal syndrome (CBS) and progressive supranuclear palsy (PSP) (1). FTD is highly heritable with 30-50% of FTD cases having a family history of the disease. Mutations in four genes including microtubule associated protein tau (*MAPT*), progranulin (*GRN*), chromosome 9 open reading frame 72 (*C9orf72*) and TANK binding kinase 1 (*TBK1*) are responsible for most familial cases (2–4).

Frontotemporal lobar degeneration (FTLD) is the pathological umbrella term used to describe FTD and similar diseases. FTLD subtypes are classified according to three proteinaceous pathological inclusions characteristic of the different diseases; tau, 43kDa transactive responsive DNA-binding protein (TDP-43) and fused in sarcoma (FUS) (5). Since the initial study by Neumann et al. (6) identified a link between hyperphosphorylated TDP-43 inclusions and FTLD, the condition, now termed FTLD with hyperphosphorylated TDP-43-positive inclusions (FTLD-TDP), has been further classified pathologically into five subtypes: A, B, C, D, and E. Across these five subtypes, the TDP-43 immunoreactive inclusions include neuronal intranuclear inclusions (NII), neuronal cytoplasmic inclusions (NCI), dystrophic neurites (DN), oligodendroglial inclusions and granulofilamentous neuronal inclusions (GFNI) (7,8). In FTLD-TDP type A pathological hallmarks consist of abundant NCI, DN and variable numbers of NII predominantly in layer II of affected cortices. Type B has high numbers of NCI in superficial and deeper cortical layers as well as the occasional DN. *C9orf72* mutations are associated with subtypes A or B. Type C is characterised by long DN which have a corkscrew appearance present in all cortical layers and is the only known subtype not linked to genetic mutations. Type D is enriched in NII, DN and occasional NCI and is associated with mutations in the *VCP* gene. Finally, Type E has GFNI present in neocortical neurons, grains and oligodendroglial inclusions.

TDP-43 is a member of the heterogeneous nuclear ribonucleoprotein (hnRNP) family and interacts with a number of other members, mainly through its C-terminal tail (9–11). The hnRNP family consists of around 20 major polypeptides, hnRNP A1-U, which range in size from 34 to 120 kDa with overlapping structural domains and common functions (12,13). HnRNPs have a key role in cellular nucleic acid metabolism, including regulating gene expression through a range of different mechanisms such as DNA maintenance and transcription, processing primary transcripts, mRNA nuclear export, subcellular localisation as well as translation and stability of mature mRNA (13–16). Perturbation of normal hnRNP activity has been observed in multiple neurological diseases, such as amyotrophic lateral sclerosis (ALS), FTLD-TDP, and Alzheimer’s disease (17–20).

Importantly, several hnRNPs have been investigated in pathological studies in relation to FTLD. HnRNP E2 accumulates in dystrophic neurites and cytoplasmic inclusions with TDP-43 in the frontal cortex and hippocampus of FTLD-TDP type A and C cases (21,22). HnRNP A3 is present in dipeptide repeat-containing NCI and NII in the hippocampus of *C9orf72* mutation carriers (23,24). HnRNP A3, H1 and H3 have also been shown to associate with hexanucleotide repeat expansions, including *C9orf72,* in disease cell and animal models (25). HnRNP K has also been shown to mislocalise from the nucleus to the cytoplasm in larger neurons, distinct from those exhibiting TDP-43 pathology in different neurodegenerative diseases, including FTLD-TDP (25,26). While their involvement in disease is increasingly recognised, these additional hnRNPs, and the network as a whole, remain significantly understudied compared to TDP-43.

Given the critical role of hnRNPs in RNA metabolism and their emerging links to neurodegeneration, this study integrates immunohistochemistry and single-nucleus RNA sequencing (snRNA-seq) to characterise hnRNP expression and localisation across three FTLD-TDP subtypes (sporadic TDP A, TDP A with *C9orf72* mutations, and TDP C). We aimed to (1) determine the diversity of TDP-43 pathology within each FTLD-TDP subtype, (2) characterise whether hnRNPs form aggregates or mislocalise across FTLD-TDP subtypes, (3) assess transcriptomic changes in hnRNP expression across cell types, and (4) evaluate concordance between transcript and protein-level alterations. Our findings indicate a lack of correlation between pTDP-43 pathology in the frontal and temporal cortices and that observed in the hippocampus. Furthermore, they suggest that the burden of hippocampal pathology may differ among distinct TDP-43 subtypes, underscoring the need to further characterise regional and subtype-specific patterns of TDP-43 pathology. Notably, several hnRNPs, including hnRNP G and Q, show cytoplasmic mislocalisation and inclusion formation, with hnRNP Q also accumulating in microglia in TDP-C. SnRNAseq and co-expression network analysis further highlights glial-enriched gene modules, particularly in astrocytes and oligodendrocytes, associated with clinical diagnosis and aggregate burden, implicating RNA splicing dysfunction as a shared molecular pathway. Together, these results underscore a critical and underappreciated glial contribution to the hnRNP network in FTLD.

## Material and Methods

### Cases

Brains were donated to Queen Square Brain Bank for Neurological Disorders, UCL Queen Square Institute of Neurology (QSBB). Cases used included FTLD-TDP type A (17 cases: TDP A 8 cases, TDP A-C9 9 cases) and TDP C (11 cases) which were diagnosed using the pathological criteria outlined by (27). This includes phosphorylated TDP-43 positive inclusions characterised by their morphology and localisation across brain regions. Our study also included neurologically normal controls which were devoid of TDP-43 pathology and had no neurological clinical manifestations (14 cases). Ethical approval for the study was obtained from the National Hospital for Neurology and Neurosurgery Local Research Ethics Committee. A demographic summary of all cases used in this study are shown in **Table 1** and extended case demographics in **Supp. Table 1**.

**Table 1:** A summary table of the case demographics for cases used in the study. Summarised are the number of cases per pathological subtype, Male to Female ratio (M/F), Age at onset, age at death and disease duration in years ± standard deviation (s.d.). Postmortem interval (PMI) is summarised as years ± standard deviation (s.d).

| **Disease Group** | **No. of cases** | **M/F** | **Age at Onset (y ± s.d.)** |
| --- | --- | --- | --- |
| TDP A | 8 | 5/3 | 65.5 ± 12.7 |
| TDP A-C9 | 9 | 5/4 | 57.2 ± 7.16 |
| TDP C | 11 | 7/4 | 60.3 ± 8.6 |
| Controls | 14 | 5/9 | n/a |

### Single nucleus RNA-seq data and analysis

The cases used in this analysis are highlighted in **Supp. Table 1**. Frozen frontal grey matter was transferred directly onto ice-cold sucrose buffer and homogenised by hand. The homogenate was layered on top of a sucrose cushion and nuclei were recovered after centrifugation. Nuclei were resuspended in PBS containing BSA (to prevent clumping) and RNase inhibitors, counted, and diluted to 1,000 nuclei/μl. Sequencing libraries were prepared using the “single cell 3’ library kit v3” with the Chromium instrument (10X Genomics). Aiming to obtain libraries for ∼5,000 independent nuclei per sample, libraries were sequenced on an Illumina NovaSeq6000 instrument (UCL Genomics), targeting an average depth of at least 100,000 reads/nucleus.

Raw sequencing data were aligned to the human reference genome (hg38) and processed through CellRanger (10X Genomics) without the inclusion of intronic reads. To assign each transcript read to its respective nucleus, we performed alignment to the transcriptome, counted reads, and processed the unique molecular identifier (UMI) tag to reduce PCR bias. The resulting gene count table was analysed using the Seurat package (v.5) (28). Quality control was performed to remove low-quality cells based on the number of detected genes and UMIs: nuclei with fewer than 200 or more than 2,500 detected genes, or with more than 5% of reads mapping to mitochondrial genes, were excluded. Cells with unusually high UMI counts, suggesting potential doublet events, were also excluded using scDblFinder (29). Using SCTransform (30), gene expression data were normalised to adjust for differences in sequencing depth between cells and scaled to correct for cell-to-cell variation in total UMI count. Mitochondrial gene content was regressed out to minimise confounding sources of variation. Batch effects were corrected using the Harmony algorithm to ensure consistency across sequencing batches. Principal component analysis (PCA) was performed on the scaled data to reduce the dimensionality of the dataset. Based on analysis of an elbow plot, the top 20 principal components were selected for downstream analyses. The PCA-generated principal components were used for clustering cells using the graph-based Louvain algorithm implemented in Seurat. Cluster annotation was performed using canonical marker genes for neural cell types sourced from brain-focused transcriptomic databases and relevant literature. (31,32). Cell types were annotated by comparing the expression pattern of marker genes within each cluster.

The snRNA-seq data was visualised using Uniform Manifold Approximation and Projection (UMAP) to view the nuclei clusters in two dimensions. Clusters were separated into the main neural cell types and differential gene expression analysis was performed to identify genes significantly differentially expressed between identified cell clusters of each disease subtype against controls using the "FindMarkers" function in Seurat with Wilcoxon rank-sum analyses, with statistical significance being defined as a false discovery rate (FDR) of < 0.05 and minimum cut-off of 3 nuclei per group.

### IPA Pathway Analysis

Significant differentially expressed genes were input into Qiagen Ingenuity Pathway Analysis software (IPA), mapped to corresponding human gene symbols from GenBank and other human gene databases (HUGO, HGNC, and Entrez gene). We then performed Core Analyses on each FTLD subtype (compared to control) across all nuclei clusters. Analyses were conducted using the human IPA Knowledge Base (Genes only) as the reference set, and all supported molecule types were included. IPA’s Upstream Regulator Analysis module predicted transcriptional regulators and signalling molecules influencing gene expression changes. Enrichment of canonical pathways and upstream regulators was assessed using a right-tailed Fisher’s exact test, with significance as defined as FDR < 0.05. Activation state predictions were based on IPA’s activation z-score algorithm with z-score > ±2 considered sufficiently dysregulated.

### hnRNP Target Jaccard Similarity Analysis

To understand the impact of TDP-43 pathology on superficial cortical neurons, we overlaid TDP-43 targets from the POSTAR3 human database: a cross-linking immunoprecipitation sequencing (CLIP-seq) database that lists proteins that preferentially bind to RBPs. The Jaccard similarity index assessed overlap in hnRNP gene targets across differentially expressed genes (DEGs) in each cell cluster, with targets retrieved from the database (33). This allowed evaluation of shared regulatory mechanisms across clusters and which neural cells exhibited the most overlap.

### hdWGCNA Analysis

High-dimensional weighted gene co-expression network analysis (hdWGCNA) was performed to assess gene co-expression modules in our dataset (34). Data were first aggregated into metacells to enhance data quality, reduce sparsity, and improve robustness of network analysis. Data were log-normalised and scaled to control for differences in sequencing depth and technical variance. Gene co-expression modules were identified with a soft-thresholding power selected based on scale-free topology fit indices. A power of 6 was chosen as it was the lowest value at which the scale-free topology model fit (R²) exceeded 0.8. Subsequently, modules were subjected to enrichment analysis across cell types within the dataset, using hypergeometric tests to identify cell type-specific gene signatures associated with each module. Analysis of relative module eigengene expression across subtypes was performed using the hdWGNCA, “FindAllDMEs” function, where modules were compared to control using a Wilcox test. These analyses allowed for the identification of disrupted gene networks associated with specific subtypes. Module-trait correlations were conducted to investigate associations between identified gene modules and clinical and pathological traits. Module eigengenes were computed as representative summaries of module expression, and Pearson correlation coefficients were calculated to quantify the strength of association between module eigengenes and traits of interest. Significant correlations were evaluated using adjusted p-values derived from two-tailed t-tests that were subjected to a Bonferroni adjustment to account for multiple hypothesis testing, thereby identifying biologically relevant modules linked to specific clinical or pathological features.

Modules from hdWGCNA showing significant associations with disease status and pathological burden were analysed for gene ontology (GO) enrichment using ClusterProfiler (35). GO terms relating to biological processes, molecular functions, and cellular components were ranked by adjusted p-values (q < 0.05), and enrichment was visualised through dot plots.

To further explore the regulatory architecture of disease-associated modules, hub gene analysis was performed within each module using hdWGCNA. Intramodular connectivity (kME) was calculated as the Pearson correlation between individual gene expression profiles and the module eigengene, representing the module’s overall expression pattern. Genes were ordered by kME values and from the top 1% were considered putative hub genes due to their strong co-expression with the core module signature. The hub genes associated with RBP mechanisms such as RNA processes, splicing, and binding were highlighted for each module network based on the overlap of related GO terms. This approach enabled the identification of central regulators potentially contributing to disease pathogenesis, particularly within astrocytic and oligodendrocyte-enriched modules that showed strong associations with disease classification and pathology scores.

The snRNA-seq data generated in this publication have been deposited in NCBI’s Gene Expression Omnibus and are accessible through GEO series accession number GSE288106.

### Immunohistochemistry for HnRNPs

Eight-micron-thick formalin-fixed paraffin-embedded (FFPE) sections were cut from TDP A cases (n=6), TDP A-C9 (n=7), TDP C (n=8) and neurologically normal control cases (n=6) (**Supp. Table 1**). The sections were deparaffinised in xylene and rehydrated using graded alcohols. Endogenous peroxidase activity was blocked using 0.3% H_2_O_2_ in methanol for 10 minutes followed by pressure cooker pretreatment for 10 minutes in citrate buffer, pH 6.0. Non-specific binding was blocked using 10% dried milk/Tris buffered saline-Tween (TBS-T) before incubating with a hnRNP derived primary antibody overnight at 4°C. **Table 2** lists all the antibodies used in this study with their supplier and concentration used. A biotinylated anti-rabbit/mouse IgG antibody (1:200, 30 minutes, DAKO) was incubated with the sections at room temperature followed by avidin-biotin complex (30 minutes, Vector Laboratories). The colour was developed with di-aminobenzidine activated with H_2_O_2_ (36). Images were taken with a Nikon H550L light microscope. Standard immunohistochemical staining for Aβ and tau were carried out to determine the presence of concomitant pathologies as described previously (37) and standard diagnostic criteria for the neuropathological diagnosis of Alzheimer’s disease and the presence of cerebral amyloid angiopathy were used in all cases (38–42).

**Table 2:** List of antibodies used in the immunohistochemical study.

| Antibody | Antibody Source | Cat.No | Specificity | Dilution | Antigen |
| --- | --- | --- | --- | --- | --- |
| TDP-43 | Cosmo | TIP-PTD-P02 | Rb | 1:10,000 | pS409/410 |
| hnRNP A1 | Abcam | ab5832 | Ms | 1:100 | Full Length |
| hnRNP A2B1 | Abcam | ab6102 | Ms | 1:100 | - |
| hnRNP C1/C2 | Abcam | ab97541 | Rb | 1:100 | aa 1-152 |
| hnRNPD1/D2 | abcam | ab61193 | Rb | 1:500 | Phospho serine 83 |
| hnRNPE1/E2 | Santa Cruz | sc-28725 | Rb | 1:100 | aa 171-280 |
| hnRNP F | abcam | ab50982 | Rb | 1:100 | aa 361-410 |
| hnRNP G | abcam | ab70064 | Rb | 1:200 | N-terminus |
| hnRNP H | abcam | ab10374 | Rb | 1:200 | aa 400-448 |
| hnRNP I | Santa Cruz | sc-16549 | Gt | 1:100 | internal sequence |
| hnRNP L | abcam | ab6106 | Ms | 1:1000 | Full length |
| hnRNP M | Sigma | HPA024344 | Rb | 1:1000 | aa 344-448 |
| hnRNP P (FUS) | Novus | NB100-565 | Rb | 1:200 | aa 1-50 |
| hnRNP Q | Thermo | PA5-15009 | Rb | 1:200 | aa 591-623 |
| hnRNP R | Abcam | ab30930 | Rb | 1:200 | - |
| hnRNP U | abcam | ab10297 | Ms | 1:1000 | Full length |

### Quantification of phosphorylated TDP-43 pathology

Quantification of the TDP-43 pathological load was undertaken in the frontal and temporal grey matter and the granule cell layer of the hippocampus. A phospho-TDP-43 (pTDP-43) antibody was used in order to stain only pathological TDP-43 inclusions that could then be quantified, without the presence of physiological TDP-43 (**Table 2**).

Since pTDP-43 is present as different pathological features in the different subtypes of FTLD, quantification of immunohistochemical staining within the frontal and temporal cortical regions was carried out manually, thereby distinguishing between intranuclear/cytoplasmic inclusions and neurites. The grid feature on QuPath software was used as a visual aid in order to ensure quantification along all six cortical layers of the grey matter per section (43). The number of inclusions and neurites were manually counted for each square within the grid and the average number of pathological features per grid square was calculated. The number of pTDP-43 inclusions were counted in the granule cell layer of the hippocampus and presented as a percentage of the total number of cells. Counting was carried out within the whole field of view at 10x magnification. Scoring of staining was performed by a single observer (AG for frontal and temporal and TL for hippocampus) blinded to clinical, histopathological and genetic status.

### Quantification of hnRNP staining

Immunohistochemically stained sections were examined microscopically to determine the cellular distribution of hnRNP staining within the frontal cortex grey matter. This region was chosen as it is involved in all subtypes of FTLD-TDP and is the region used in the gene expression analysis. The degree of neuronal nuclear and/or cytoplasmic hnRNP immunostaining was scored semi-quantitatively as previously described (22,24). All the grey matter within a tissue section was analysed, and the following rating scale was employed: 0 = no staining present; 1 = few (1–5) cells showing weak nuclear and/or cytoplasmic staining; 2 = moderate number (5–10) cells showing moderate nuclear and/or cytoplasmic staining; 3 = more than 10 cells showing strong nuclear and/or cytoplasmic staining. The severity of hnRNP immunostaining found in pathological inclusions (ie neuronal cytoplasmic, neuronal intranuclear and dystrophic neurites) in the grey matter of the frontal cortex was also graded employing the following rating scale: 0 = no inclusions present; 0.5 = rare (ie 1-5 inclusions per section); 1 = few (ie 1-5 inclusions per field); 2 = mild (ie 5-10 inclusions per field); 3 = moderate (ie 10-50 inclusions per field). Any variation to the normal hnRNP staining pattern was also documented. Scoring of staining was performed by a single observer (TL) blinded to clinical, histopathological and genetic status.

### Statistical analysis

Group comparisons of age at onset, age at death, *post-mortem* interval and duration of illness were made using Kruskal-Wallis test with post-hoc Dunn’s multiple comparison test. pTDP-43 immunostaining data was analysed using GraphPad Prism software (version 10.0). The 28 FTLD patients were stratified according to genetic and pathological subtype for statistical analysis of the effect of each mutation and underlying pathology on the degree and pattern of pTDP-43 staining. Comparisons of semi-quantitative scores for the frequency of pTDP-43 positive inclusions and neurites in neurons of the frontal and temporal cortex, and dentate gyrus of the hippocampus, with respect to pathological type or genetic mutation, were performed using Kruskal-Wallis test with post-hoc Dunn’s multiple comparison test.

Comparison of rating scores for the inclusion/cytoplasmic/nuclear (cellular localisation) immunostaining of the various hnRNPs in the three different pathological subtypes were performed using a Kruskal-Wallis H test (K independent samples) on SPSS software (v.29), a non-parametric method appropriate for comparing ordinal data between more than two independent groups. Specifically, separate Kruskal-Wallis tests were performed to assess differences in protein scores (scored 0–3) across the four disease conditions (Control, TDP A, TDP A-C9, TDP C) within each cellular compartment (nucleus, cytoplasm, inclusion). The HnRNP cellular localisation was deemed as the test variable whilst the pathological subtype was the grouping variable. The H statistic for this test reflects the degree to which the mean ranks differ among groups; a higher H value indicates greater differences between groups. In all instances, significance levels for the p value were set at p < 0.05.

## Results

### Demographic comparisons

The demographic data pertaining to the cases included in this study are listed in **Table 1**. There was a statistical difference in *post-mortem* interval between the FTLD subtypes and control groups (H=9.979, p=0.019) though pairwise comparisons revealed no statistically significant difference. There was no statistical difference between age of onset (H=2.505, p=0.286) and disease duration (H=4.606, p=0.099) in the FTLD subtype groups (**Supp. Fig. 1**). The mean age at death (AAD) was statistically different between the groups (H=9.632, p=0.022) with the TDP A-C9 group having a statistically lower mean AAD than controls (Dunn’s multiple comparison, adjusted p=0.012; **Supp. Fig. 1**).

### Variation in pTDP-43 pathology burden across and within FTLD pathological subtypes

Cases from the FTLD pathological subtypes Type A (TDP A sporadic and genetic) and Type C (TDP C) exhibited the classic pTDP-43 pathological features of neuronal cytoplasmic inclusions (NCI), short thick dystrophic neurites (DN), lentiform neuronal intranuclear inclusions (NII) (TDP A), and long thick DN (TDP C), respectively (**Fig. 1**). The abundance of these features, however, varied case by case as visible in the pathology scoring in **Supp. Table 1**. Manual quantification of neurites (long or short DN) and inclusions (NCI and NII) was therefore carried out to generate a semi-quantitative assessment of pathology burden per case. Combined pathology burden (inclusions and neurites) was greatest in the TDP C cases within the frontal cortex (Kruskal Wallis test H=10.39, p=0.006, Dunn’s multiple comparison test, **Fig. 1 a, b**). TDP A-C9 exhibited the lowest amount of combined pathology in both the frontal and temporal cortex (**Fig. 1 b, c**). This was mainly attributable to the lack of pTDP-43 positive neurites in these cases. TDP A sporadic cases exhibited the greatest amount of pathological inclusions within the frontal cortex (Kruskal Wallis test H=6.142, p=0.046272, Dunn’s multiple comparison test, **Fig. 1 d**) As expected, TDP C cases exhibited a low amount of inclusions and the greatest amount of neurites within the frontal cortex (Kruskal Wallis test H= 13.33, p=0.001, Dunn’s multiple comparison test, **Fig. 1 d, e**). Ultimately, pTDP-43 pathology was greater in the frontal cortex than in the temporal cortex but there was an overall strong correlation between the two regions (Simple Linear regression, R squared = 0.349, p= 0.002, **Fig. 1 f**).

**Fig. 1.**
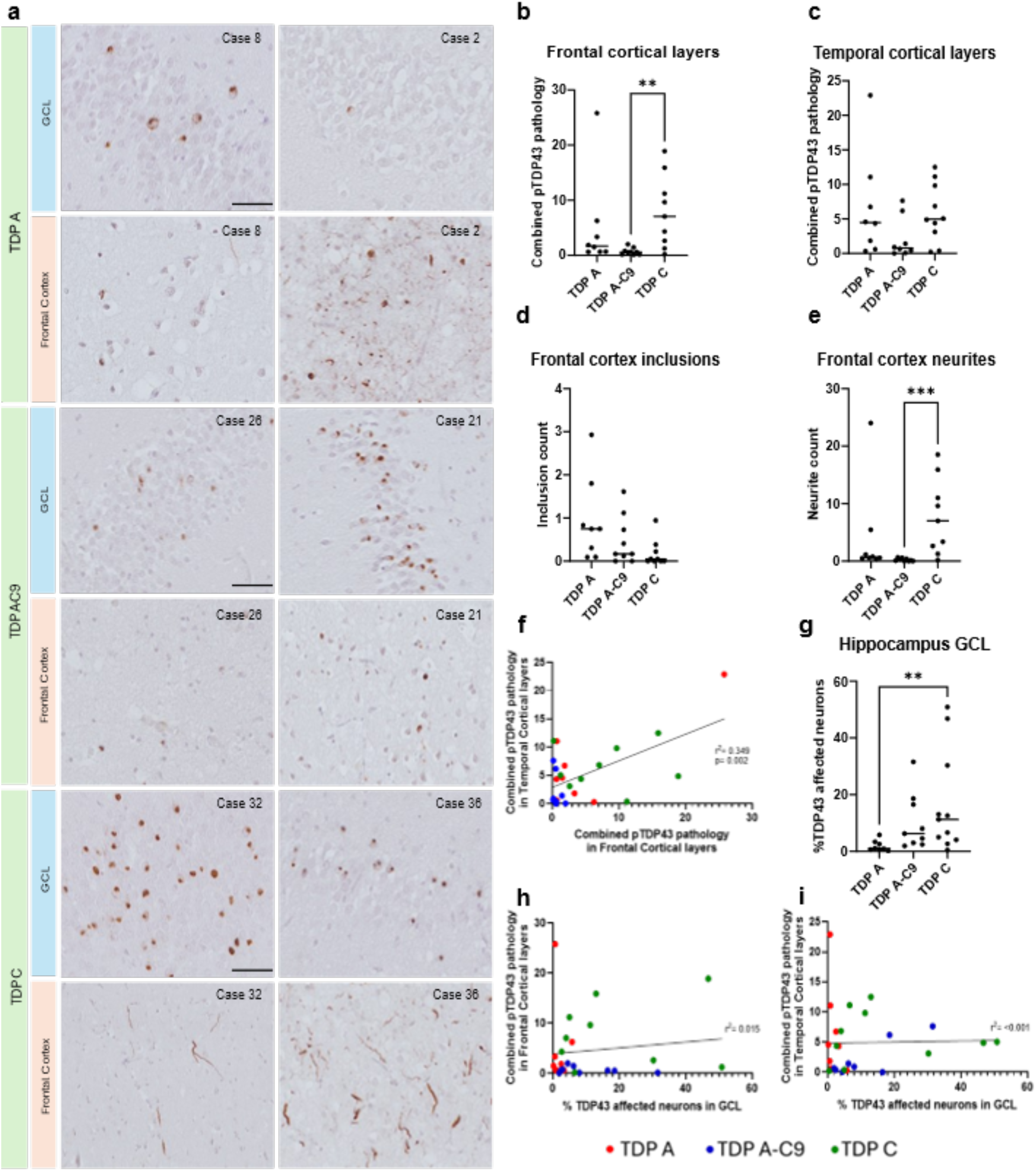
Quantification of phosphorylated TDP-43 (pTDP43) pathology in FTLD-TDP subtypes. (a) Immunohistochemical images showing that frequency of pTDP-43 pathological inclusions and/or neurites varies greatly between cases and across subtypes. (b) Quantification of combined pTDP43 pathology (inclusions and neurites) in the frontal cortical layers across subtypes TDP A (n=8), TDP A-C9 (n=9), and TDP C (n=11). (c) Quantification of combined pTDP43 pathology (inclusions and neurites) in the temporal cortical layers across subtypes TDP A, TDP A-C9 and TDP C. (d) Quantification of pTDP43 positive inclusions in the frontal cortical layers across subtypes TDP A, TDP A-C9 and TDP C. (e) Quantification of pTDP43 positive neurites in the frontal cortical layers across subtypes TDP A, TDP A-C9 and TDP C. (f) Correlation of combined pTDP43 pathology along temporal and frontal cortical layers (Simple Linear regression, R squared = 0.349, p= 0.002). (g) Quantification of pTDP43 pathology in the hippocampal GCL. Values are shown as percentage of cells affected by pTDP43 inclusions in relation to total number of cells. (h) Correlation of combined pTDP43 pathology along GCL in hippocampus and frontal cortical layers (Simple Linear regression, R squared = 0.015, p= n.s.). (i) Correlation of combined pTDP43 pathology along GCL in hippocampus and temporal cortical layers (Simple Linear regression, R squared = <0.001, p= n.s.). All comparisons of pathology between FTLD subtypes were analysed by Kruskal-Wallis tests with Dunn’s multiple comparison tests. In relation to all statistical results, *=p=0.01 to 0.05, **=p=0.001 to 0.01 and ***=p=<0.001. A p value > 0.05 is deemed n.s. (non-significant). Demographic data of the cases outlined in this figure are listed in Supp. Table 1. Scale bar represents 50µm.

pTDP-43 positive neuronal cytoplasmic inclusions were also common within the granular cell layer (GCL) of the dentate gyrus in the hippocampus in FTLD-TDP (44,45). We therefore sought to quantify TDP-43 pathology in the GCL within the same cases. TDP C cases exhibited the greatest amount of pTDP-43 affected neurons within the GCL (Kruskal Wallis test H=9.526, p=0.009, Dunn’s multiple comparison test, **Fig. 1 a, g**). Interestingly, of the TDP A subgroup, the *C9orf72* mutant cases (TDP A-C9) exhibited the greatest TDP-43 pathology burden in the GCL, paradoxical to what is observed in the neocortical regions (**Fig. 1 a, g**). Of note, pTDP-43 pathology within the hippocampal GCL does not correlate to TDP-43 pathological burden in the frontal or temporal regions within the same cases (**Fig. 1 h, i**).

Importantly, the variability in frequency of pTDP-43 pathology is clear, even amongst cases within the same pathological subgroup. Cases with similar disease duration and age of onset can exhibit great variability in the amount of pTDP-43 pathology in both frontal/temporal cortical regions and the hippocampus (**Supp. Table 1)**.

### HnRNP localisation is disrupted and can lead to pathological inclusions in FTLD-TDP

Since TDP-43, a protein belonging to the hnRNP family, showed variable levels of pathology in these cases, we chose to investigate the expression and cellular localisation of additional hnRNPs. We therefore chose to carry out a semi-quantitative pathological assessment of multiple hnRNPs in the different FTLD subtypes and controls. We investigated the presence and localisation of the hnRNPs in the cortical layers of the frontal cortex, focusing mainly on immunohistochemical staining within neurons (**Supp. Fig. 2-4**). The median scores (with interquartile range) from hnRNP immunostaining of pathological inclusions, as well as a staining score within neuronal nuclei and cytoplasm, are shown in **Table 3**.

**Table 3:** Semi-quantification of the immunohistochemical analysis of the hnRNPs in different FTLD subtypes. Values are listed in the range 0-3 where 0 = staining is absent, 1 = few cells stained, 2= moderate number of cells stained and 3 = many cells stained. Differences in group distribution are statistically analysed via a Kruskal-Wallis K independent sample test. H indicates the H-test statistic and the p value indicates statistical significance. Statistically significant tests where p<0.05 are written in bold suggesting there is a significant altered distribution of the HnRNP in an FTLD subtype compared to controls.

| HnRNP A1 |  |  |  | HnRNP A2B1 |  |  |
| --- | --- | --- | --- | --- | --- | --- |
|  | Inclusions | Nucleus | Cytoplasm | Inclusions | Nucleus | Cytoplasm |
| TDP A | 0 (0-0) | 2 (1-2) | 1 (1-2) | 0 (0-0) | 3 (3-3) | 1 (1-2) |
| TDP A-C9 | 0 (0-0) | 2 (2-3) | 2 (1-2) | 0 (0-0) | 3 (2-3) | 2 (1-2) |
| TDP C | 0 (0-0) | 2 (2-3) | 2 (1-2) | 0 (0-0) | 3 (2-3) | 2 (1-2) |
| Controls | 0 (0-0) | 3 (3-3) | 2 (2-2) | 0 (0-0) | 3 (3-3) | 0 (0-1) |
|  | H=0 | H=13.791 | H=9.913 | H=0 | H=3.468 | H=17.501 |
|  | p=1.00 | p=0.003 | p=0.019 | p=1.00 | p=0.325 | p=<0.001 |
| HnRNP C |  |  |  | HnRNP D |  |  |
|  | Inclusions | Nucleus | Cytoplasm | Inclusions | Nucleus | Cytoplasm |
| TDP A | 0 (0-0) | 3 (3-3) | 0 (0-0) | 0 (0-0) | 2 (2-3) | 0 (0-1) |
| TDP A-C9 | 0 (0-0) | 3 (3-3) | 0 (0-0) | 0 (0-0) | 2 (2-3) | 1 (0-1) |
| TDP C | 0 (0-0) | 3 (3-3) | 0 (0-0) | 0 (0-0) | 3 (3-3) | 2 (1-2) |
| Controls | 0 (0-0) | 3 (3-3) | 1 (0-1) | 0 (0-0) | 3 (3-3) | 0 (0-0) |
|  | H=0 | H=0 | H=15.826 | H=0 | H=11.243 | H=18.614 |
|  | p=1.00 | p=1.00 | p=0.001 | p=1.00 | p=0.01 | p=<0.001 |
| HnRNP E1/2 |  |  |  | HnRNP F |  |  |
|  | Inclusions | Nucleus | Cytoplasm | Inclusions | Nucleus | Cytoplasm |
| TDP A | 0 (0-0) | 0 (0-0) | 0 (0-2) | 0 (0-0) | 0 (0-1) | 3 (3-3) |
| TDP A-C9 | 0 (0-0) | 0 (0-0) | 0 (0-1) | 0 (0-0) | 1 (0-1) | 3 (3-3) |
| TDP C | 1 (0-1) | 2 (1-2) | 1 (1-2) | 0 (0-0) | 0 (0-1) | 2.5 (2-3) |
| Controls | 0(0-0) | 0(0-0) | 1 (1-1) | 0 (0-0) | 0 (0-0) | 3 (3-3) |
|  | H=16.156 | H=24.504 | H=9.096 | H=0 | H=5.574 | H=10.739 |
|  | p=0.001 | p=<0.001 | p=0.028 | p=1.00 | p=0.134 | p=0.013 |
| HnRNP G |  |  |  | HnRNP H |  |  |
|  | Inclusions | Nucleus | Cytoplasm | Inclusions | Nucleus | Cytoplasm |
| TDP A | 1 (1-1) | 1 (1-1) | 1 (1-1) | 0 (0-0) | 2 (2-3) | 0.5 (0-2) |
| TDP A-C9 | 1 (1-1) | 1 (1-2) | 1 (1-1) | 0 (0-0) | 2 (2-3) | 1.5 (1-2) |
| TDP C | 1 (0-1) | 1 (1-1) | 1 (1-2) | 0 (0-0) | 3 (2-3) | 2 (2-3) |
| Controls | 0 (0-0) | 1 (1-1) | 2 (2-2) | 0 (0-0) | 3 (3-3) | 0 (0-0) |
|  | H=18.552 | H=2.714 | H=15.097 | H=0 | H=13.004 | H=17.056 |
|  | p=<0.001 | p=0.438 | p=0.002 | p=1.00 | p=0.005 | p=<0.001 |
| HnRNP I |  |  |  | HnRNP L |  |  |
|  | Inclusions | Nucleus | Cytoplasm | Inclusions | Nucleus | Cytoplasm |
| TDP A | 0 (0-0) | 2 (2-2) | 1 (0-2) | 0 (0-0) | 3 (3-3) | 1 (0-1) |
| TDP A-C9 | 0 (0-0) | 2 (2-2) | 1 (0-1) | 0 (0-0) | 3 (3-3) | 1 (0-3) |
| TDP C | 0 (0-0) | 2 (2-3) | 1 (1-2) | 0 (0-0) | 3 (3-3) | 2 (1-3) |
| Controls | 0 (0-0) | 3 (3-3) | 0.5 (0-1) | 0 (0-0) | 3 (3-3) | 2 (1-2) |
|  | H=0 | H=17.717 | H=9.673 | H=0 | H=0 | H=7.786 |
|  | p=1.00 | p=<0.001 | p=0.22 | p=1.00 | p=1.00 | P=0.051 |
| HnRNP M |  |  |  | HnRNP P (FUS) |  |  |
|  | Inclusions | Nucleus | Cytoplasm | Inclusions | Nucleus | Cytoplasm |
| TDP A | 0 (0-0) | 3 (3-3) | 0 (0-0) | 0 (0-0) | 3 (2-3) | 0 (0-0) |
| TDP A-C9 | 0 (0-0) | 3 (3-3) | 0 (0-0) | 0 (0-0) | 3 (3-3) | 0 (0-1) |
| TDP C | 0 (0-0) | 3 (3-3) | 0 (0-0) | 0 (0-0) | 3 (3-3) | 1 (1-2) |
| Controls | 0 (0-0) | 3 (3-3) | 0 (0-0) | 0 (0-0) | 3 (3-3) | 2 (1-2) |
|  | H=0 | H=0 | H=0 | H=0 | H=4.000 | H=19.284 |
|  | p=1.00 | p=1.00 | p=1.00 | p=1.00 | p=0.261 | p=<0.001 |
| HnRNP Q |  |  |  | HnRNP R |  |  |
|  | Inclusions | Nucleus | Cytoplasm | Inclusions | Nucleus | Cytoplasm |
| TDP A | 0.5 (0-1) | 1.5 (1-3) | 1 (1-2) | 0 (0-1) | 2.5 (0-3) | 2 (0-3) |
| TDP A-C9 | 1 (0-1) | 2 (1-3) | 1 (1-2) | 0 (0-1) | 2 (0-3) | 1.5 (0-3) |
| TDP C | 1 (0-1) | 3 (2-3) | 2 (1-2) | 0 (0-1) | 2 (2-3) | 2 (2-3) |
| Controls | 0 (0-0) | 2.5 (2-3) | 1.5 (1-2) | 0 (0-0) | 2.5 (1-3) | 2 (1-2) |
|  | H=5.584 | H=10.422 | H=7.405 | H=1.962 | H=0.807 | H=1.629 |
|  | p=0.134 | p=0.015 | p=0.06 | p=0.580 | p=0.848 | p=0.653 |
| HnRNP U |  |  |  |  |  |  |
|  | Inclusions | Nucleus | Cytoplasm |  |  |  |
| TDP A |  |  |  |  |  |  |
| TDP A-C9 | 0 (0-0) | 2.5 (1-3) | 0 (0-0) |  |  |  |
| TDP C | 0 (0-0) | 2 (2-3) | 0 (0-0) |  |  |  |
| Controls | 0 (0-0) | 2 (2-3) | 0 (0-0) |  |  |  |
|  | 0 (0-0) | 2 (2-2) | 0 (0-0) |  |  |  |
|  | H=0 | H=2.147 | H=0 |  |  |  |
|  | p=1.00 | p=0.542 | p=1.00 |  |  |  |

In control cases, hnRNP proteins A2/B1, D1/2, H1, and I were predominantly localised to the nucleus, with minimal cytoplasmic presence **(Supp. Fig. 2,3)**. However, in FTLD cases, these hnRNP proteins showed increased cytoplasmic localisation across all pathological subtypes, indicating a shift from their usual nuclear distribution. Contrastingly, hnRNP E1/E2 and F were primarily cytoplasmic in controls, suggesting a more physiological role in the cytosol under normal conditions (**Supp. Fig. 2,3**). While hnRNP F exhibited heightened nuclear staining in disease subtypes compared to controls, hnRNP M and U maintained their nuclear localisation across both control and FTLD cases, with no observed alterations in cellular distribution **(Supp. Fig. 4)**. HnRNP C also exhibited a predominant nuclear presence in both the controls and FTLD cases, but there was a cytoplasmic presence observed in the controls that was not seen in the FTLD cases **(Supp. Fig. 2).** HnRNP A1, L, G, R and Q were all observed in both the nucleus and cytoplasm and this distribution was not altered in disease. Importantly, no significant differences in hnRNP localisation or staining intensity were noted between TDP A (sporadic) and TDP A-C9 cases (**Supp. Fig. 2-4**).

Pathological inclusions of hnRNP proteins were observed in various cell types across FTLD-TDP subtypes, showing distinct patterns of accumulation that may contribute to disease pathology. HnRNP E1/E2 showed distinct expression in dystrophic neurites in TDP C cases, consistent with previous studies (21,22) (**Fig. 2a-d**). HnRNP G accumulation was particularly prominent in non-neuronal cells, showing consistent accumulation in astrocytes across all FTLD subtypes (**Fig. 2e-h**). HnRNP G astrocytic morphologies were present throughout the cortical layers and varied from accumulation within the astrocytic cell bodies to the processes. HnRNP R, on the other hand, showed distinct patterns of accumulation in NCIs and axonal projections, as seen in sporadic TDP A (**Fig. 2i**) and TDP A-C9 (**Fig. 2j**). In TDP C, hnRNP R was present in axonal projections (**Fig. 2k**), indicating a possible involvement in axonal transport or structural integrity within affected neurons. The accumulation patterns of hnRNP Q were notably varied across FTLD subtypes, indicating a complex involvement in disease pathology. HnRNP Q intensely stained astrocytes in both sporadic TDP A (**Fig. 2l-m**) and TDP C cases (**Fig. 2q**), highlighting its strong association with astrocytic pathology in FTLD. In contrast, in TDP A-C9 cases, hnRNP Q was primarily seen in neuritic profiles, suggesting subtype-specific distribution in neuronal projections (**Fig. 2n**). HnRNP Q was also observed to accumulate in microglia in TDP C (**Fig. 2o-p**), suggesting a subtype-specific involvement in glial cell types, potentially linked to inflammatory or neurodegenerative processes within these cells. As hnRNP Q accumulation was observed in both astrocytes and microglia in TDP C cases, it may highlight a broader role in glial pathology within this subtype.

**Fig. 2:**
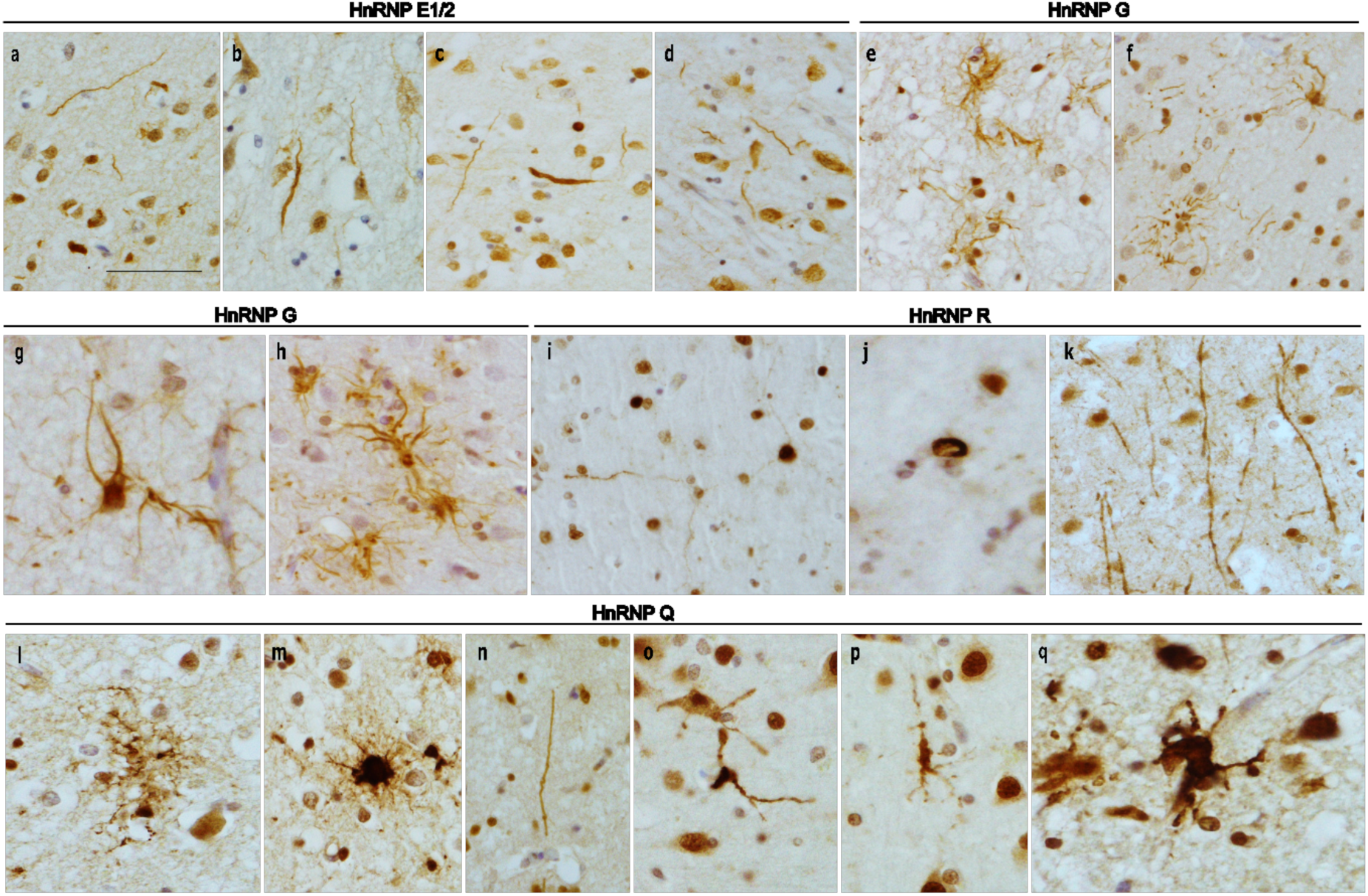
HnRNP pathologies present in FTLD-TDP subtypes. HnRNP E is observed in neurites in FTLD TDP C (a and b, case 34; c and d, case 29). HnRNP G accumulation is seen in astrocytes in all subtypes, sporadic TDP A (e, case 1), TDP A-C9 (f, case 20) and TDP C (g, case 29 and h, case 32). HnRNP R accumulation is present in neuronal cytoplasmic inclusions and neuronal processes in sporadic TDP A (i, case 5), cytoplasmic inclusions in TDP A-C9 (j, case 26) and in axonal projections in TDP C (k, case 34). HnRNP Q accumulation is seen in all FTLD-TDP subtypes although different cellular morphologies are affected. In sporadic TDP A astrocytic profiles show an accumulation of hnRNP Q (l and m, case 1), in TDP A-C9. HnRNP Q is seen in neuritic profiles (n, case 19). In TDP C cases both microglia (o and p, case 34) and astrocytes (q, case 32) are seen to accumulate hnRNP Q. Bar in a represents 50µm in a-f, h, i, k and 30µm in g, j, l-q.

### Single-nuclei transcriptomic distribution of FTLD-TDP subtypes compared to control

To determine how each cell type in the frontal cortex was affected in FTLD-TDP, we undertook a snRNA-seq study in a subset of sporadic TDP A (n=3), TDP A-C9 (n=3) cases and TDP C (n=6), along with neurologically normal controls without TDP-43 pathology (n=5; **Fig. 3a, Supp. Table 1**). After quality control preprocessing and excluding two samples for not passing checks (two TDP C cases), we recovered a total of 93,833 nuclei across all samples of which we identified a total of 34 clusters (**Fig. 3b i and ii, Supp. Table 2**). Within the 34 clusters sequenced, we identified all the main neural cell types (**Fig. 3c**). This included 7 excitatory neuronal clusters, 5 distinct inhibitory neuronal clusters (*VIP, SST, PVALB, LAMP5*, and *RELN* expressing), as well as glial cells; 6 oligodendrocyte, 2 microglial, 4 astrocytic, and 2 oligodendrocyte precursor cell (OPCs) clusters (**Supp. Table 2**). We also collected nuclei from endothelial cells, fibroblasts, and pericytes, as well as other cells relating to vasculature (peripheral immune cells, leptomeningeal cells). We began by inferring cortical layer specificity of excitatory neurons through comparative mapping to annotated reference datasets from the CZ CellxGene Discover database and relevant published snRNA-seq datasets. This enabled classification of excitatory neuronal subtypes based on their transcriptomic alignment with known laminar markers. Following this, we identified significant changes in differential gene expression in the L2-3 excitatory neurons, mature oligodendrocytes, OPCs, and astrocytes across all FTLD subtypes compared to control, with more significant changes seen in L3-5 in sporadic TDP A and TDP C (**Fig. 3d i and iii**).

**Fig. 3:**
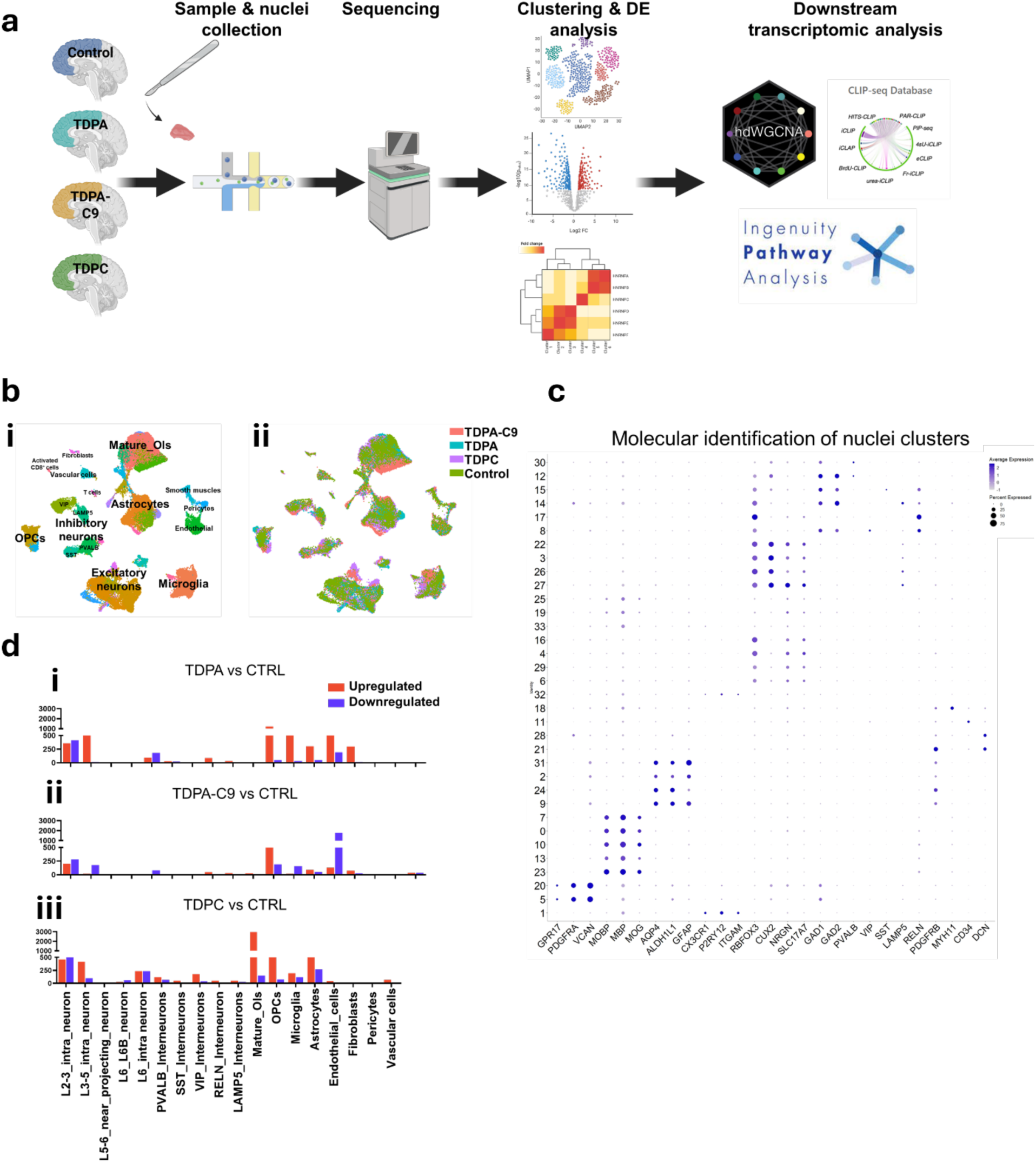
Workflow and characterisation of cell types and molecular profiles in FTLD subtypes: (a) Schematic representation of the experimental workflow. Frontal cortex samples from FTLD-TDP subtypes (TDP A; n=3, TDP A-C9; n=3, and TDP C; n=4) and normal controls (NC; n=5) were dissociated into single nuclei, followed by single-nucleus RNA sequencing (snRNA-seq) and downstream computational analysis. (b) UMAP plots displaying the clustering of 93,833 single nuclei, (i) annotated into major cell types, including astrocytes, microglia, excitatory neurons, inhibitory neurons, mature oligodendrocytes, oligodendrocyte progenitor cells (OPCs), and vascular cells and highlighting nuclei distribution across FTLD subtypes (ii; TDP A, TDP C, TDP A-C9) and normal controls. (c) Dot plot summarising the molecular characterisation of numbered clusters from 0-32. Dot size represents the percentage of cells expressing the marker gene, and colour intensity indicates average expression level. (d) Bar chart indicating total number of significant differentially expressed genes across all neural cell clusters of (i) TDPA, (ii) TDPA-C9, and (iii) TDPC compared with control. Graphs represent positive and negative log₂ fold changes as red and blue, respectively, with significance being set at FDR <0.05 using a Wilcox test followed by the Benjamini-Hochberg method.

### Transcriptomic impact on excitatory neurons across FTLD-TDP subtypes

We investigated L2-3 excitatory neurons in the frontal cortex, due to their higher vulnerability to TDP-43 pathology in FTLD-TDP (**Fig. 4a i**). While all three FTLD subtypes share a core set of DEGs (**Fig. 4a ii, Supp. Table 2**), TDP A and TDP A-C9 were more transcriptionally similar, with overlapping yet subtype-enriched profiles (**Supp. Fig. 5**). TDP C exhibits a largely distinct transcriptomic landscape, as evidenced by its separation in dendrogram clustering. It also showed the highest number of unique DEGs. While this may reflect genuine subtype-specific mechanisms of neuronal vulnerability and dysfunction in FTLD, we acknowledge that the larger sample size and greater number of cells for TDP C may have contributed to increased statistical power and DEG detection. As such, caution is warranted when interpreting DEG counts across conditions or cell types, as they may not represent equivalent comparisons. (**Fig. 4a ii, Supp. Fig. 5**). We therefore performed canonical pathway analysis on L2-3 excitatory neurons in the frontal cortex to identify inferred functional alterations across different FTLD subtypes compared to controls (**Fig. 4a iii, Supp. Table 3**). In the TDP A subgroup, a consistent upregulation of pathways involved in translational control and protein metabolism was observed. These included EIF2AK4 (GCN2) response to amino acid deficiency, Eukaryotic Translation Elongation, Translation Initiation and Termination, and Nonsense-Mediated Decay (NMD), all showing significant negative regulation, indicative of downregulation. Signalling by ROBO receptors was the only pathway upregulated in this subtype, suggesting potential alterations in axon guidance mechanisms. Additional downregulated pathways involved SRP-dependent co-translational protein targeting, selenocysteine metabolism, and rRNA processing, pointing toward broader deficits in proteostasis and ribosomal function. The TDP A-C9 subgroup showed partial transcriptional overlap with sporadic TDP A (**Supp. Fig. 5**), particularly in the downregulation of EIF2AK4 signalling, translation-related pathways, and NMD. However, this subtype also exhibited a uniquely upregulated Synaptogenesis Signalling Pathway, suggesting that aberrant synaptic remodelling may be a distinguishing molecular feature in *C9orf72*-associated FTLD. Other pathways such as mitochondrial RNA degradation and major rRNA processing remained downregulated, reinforcing the disruption of RNA metabolism in these neurons. In TDP C, the pathway profile closely mirrored that of TDP A, with robust downregulation of translation-related pathways, including EIF2AK4 response, elongation/initiation/termination, and NMD. Like TDP A, ROBO signalling appeared as upregulated in TDP C, indicating a convergent upregulation of axonal guidance cues across sporadic subtypes. Unique to TDP C was a pronounced downregulation in selenocysteine metabolism, suggesting metabolic vulnerabilities specific to this variant.

**Fig. 4.**
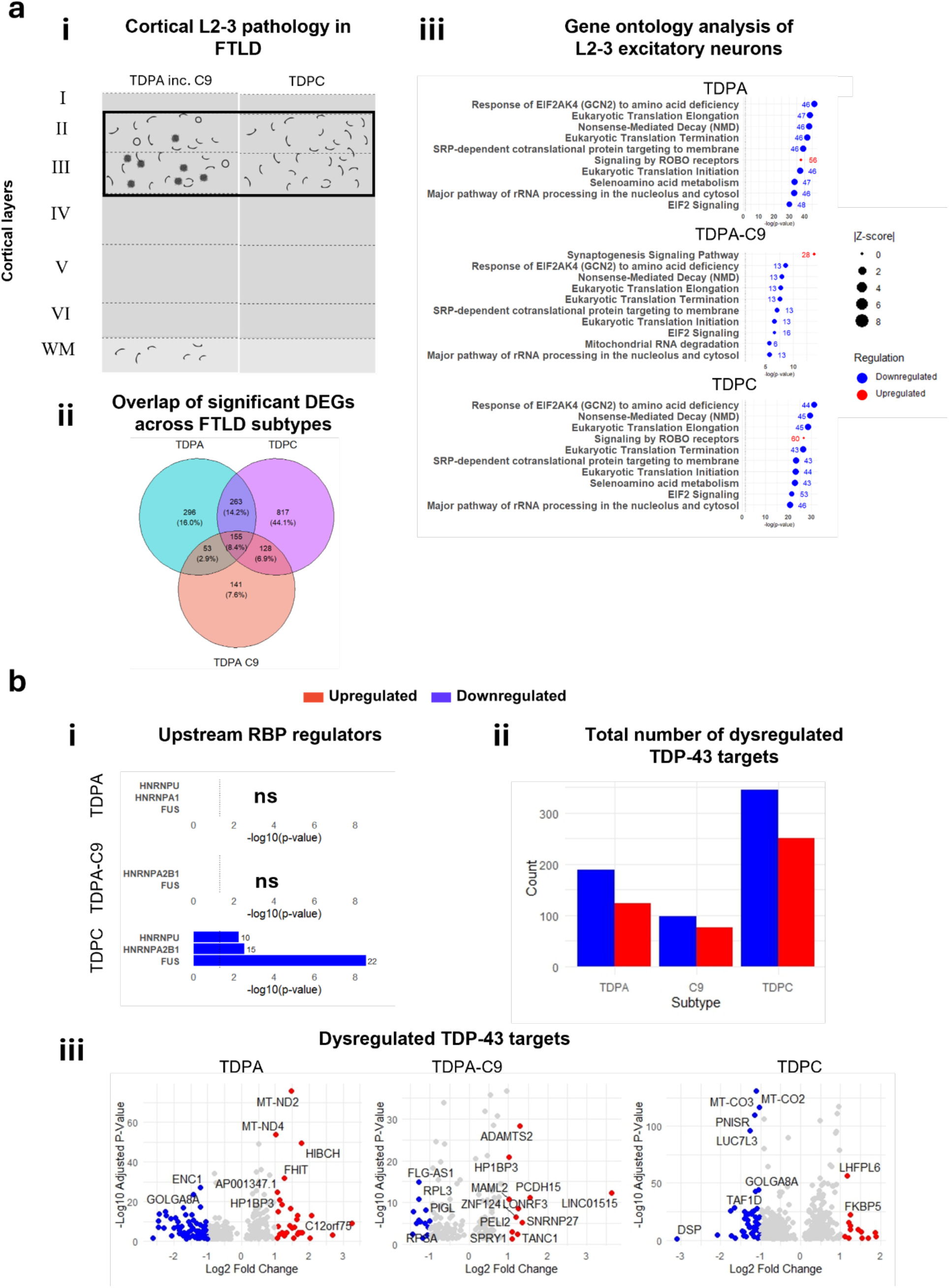
Characterisation of L2-3 excitatory neurons across FTLD-TDP subtypes: (a) (i) Schematic illustration of TDP-43 pathology in the cortex of FTLD-TDP subtype A (TDPA; n=3 and TDPA-C9; n=3) and subtype C (TDPC; n=4) cases, with a focus on L2-3 cortical layers. (ii) Venn diagram showing the overlap of significantly differentially expressed genes (DEGs) between FTLD-TDP subtypes relative to control in L2-3 excitatory neurons. Total gene counts are indicated, with the percentage of the total DEG pool shown in parentheses. (iii) Dot plot of the top 10 canonical pathways enriched in L2-3 excitatory neurons across FTLD-TDP subtypes compared to control. Pathway significance was determined by right-tailed Fisher’s exact test, with a -log₁₀ p-value threshold of 1.3 (dotted line). Dot size reflects the predicted activation z-score, while colour denotes activation status (red = positive, blue = negative). The number adjacent to each dot indicates the number of DEGs associated with that pathway. (b) (i) Upstream regulator analysis focused on heterogeneous nuclear ribonucleoproteins (hnRNPs) across FTLD-TDP subtypes. Regulators are ranked by -log₁₀ adjusted p-value, with direction of enrichment (up- or down-regulated) highlighted. NS = not significant. (ii) Bar plot showing the total number of DEGs overlapping with POSTAR3-derived TDP-43 targets, stratified by expression direction (red = upregulated, blue = downregulated). (iii) Volcano plots of differentially expressed TDP-43 target genes in each FTLD subtype (TDPA, TDPC, and TDPA with *C9orf72* mutation) relative to controls. Red and blue dots indicate significantly up- or downregulated genes, respectively, based on -log₁₀ adjusted p-value and log₂ fold change. Selected genes of interest are labelled.

Despite the well-established dysfunction of TDP-43 in FTLD-TDP, *TARDBP* itself was not differentially expressed in L2-3 neurons compared to controls. To explore the downstream consequences of altered TDP-43 function, we performed upstream regulator analysis focused on RBPs, using the differentially expressed genes from the L2-3 neuronal cluster to predict regulatory disruptions at the post-transcriptional level (**Fig. 4b i**). Whilst we did not report any predicted changes in TDP-43, we identified disrupted regulation of other RBPs including HNRNPA2B1, HNRNPU, and FUS. These RBPs were present across all subtypes, however, TDP C was the only subtype that exhibited significant predicted changes. To further confirm alterations in TDP-43 functionality, we investigated the dysregulated POSTAR3 TDP-43 targets, which revealed a marked dysregulation of targets, mostly in TDP C and, to a lesser extent, TDP A cases, supporting widespread post-transcriptional dysregulation (**Fig. 4b ii)**. TDP A-C9 cases showed fewer total dysregulated targets in this layer, with a modest skew toward downregulation. At the gene level, TDP A samples exhibited mitochondrial transcript upregulation (*MT-ND2*, *MT-ND4*), and TDP C showed downregulation of genes such as *MT-CO2* and *PNISR*, many of which are known or predicted TDP-43 targets (**Fig. 4b iii**). In L3-5 neurons, where TDP-43 pathology was less abundant (**Fig. 5a i**), the transcriptional changes in FTLD were more fragmented across subtypes, with very few DEGs shared across all (**Fig. 5a ii**). Sporadic TDP A appears highly distinct and shows minimal overlap with TDP A-C9. In contrast, TDP A-C9 and TDP C share a larger DEG subset, suggesting greater transcriptional similarity in this layer (**Supp. Fig. 5**). Pathway analysis of L3-5 excitatory neurons revealed pronounced alterations in synaptic signalling and neurovascular pathways across FTLD subtypes (**Fig. 5a iii, Supp. Table 3**). In TDP A cases, there was a marked upregulation of pathways governing synaptic structure and neurotransmission, including synaptogenesis signalling, endocannabinoid neuronal synapse, and both glutamatergic and GABAergic receptor signalling. Additional enrichment of Neurexin/Neuroligin and Netrin signalling pathways pointed to potential dysregulation of axonal connectivity and excitatory-inhibitory balance. Notably, neurovascular coupling signalling was upregulated, suggesting broader vascular and stress-related responses in these deeper-layer neurons. TDP A-C9 neurons showed a distinct signature dominated by downregulation of mitochondrial and bioenergetic pathways, including mitochondrial RNA degradation, oxidative phosphorylation, respiratory electron transport, and complex I biogenesis. This was accompanied by reduced activity in tRNA processing and mitochondrial protein degradation, implicating widespread impairments in mitochondrial quality control and energetic homeostasis. TDP C samples revealed a robust enrichment of synaptic and signalling-related pathways, similar to TDP A, but with notable subtype-specific differences. Upregulation of synaptic long-term potentiation and SNARE signalling pathways indicated alterations in vesicle trafficking and plasticity. As with TDP A, opioid and glutamatergic receptor signalling pathways were prominently elevated, reinforcing excitatory-inhibitory imbalance as a convergent feature of non-C9 FTLD. Further analysis showed stronger enrichment of RBPs as upstream regulators in L3-5 compared to L2-3 (**Fig. 5b i)**. More RBP regulators such as *HNRNPA2B1*, *HNRNPU*, and *TARDBP* reached significance, and their relative statistical enrichment was higher and often subtype-specific. For example, HNRNPR and FUS were most prominent in TDP A cases, while TARDBP and HNRNPU appeared in both TDP A and TDP A-C9. The downstream impact on TDP-43 targets was also more variable: TDP A showed the greatest number of dysregulated targets, again skewed toward upregulation, while TDP A-C9 showed lower enrichment for TDP-43 targets, and, in contrast to L2-3 neurons, TDP C retained a moderate upregulation profile (**Fig. 5b ii**). Volcano plots of TDP-43 targets from TDP A L3-5 neurons, such as *EFNA5* and *FRAS1,* were upregulated, pointing to altered axon guidance. TDP A-C9 displayed downregulation of mitochondrial genes (e.g., *MT-CO2*) and *TARBP1*, while TDP C neurons upregulated *CALM1*, *NRGN*, and *ENC1* (**Fig. 5b iii**).

**Fig. 5.**
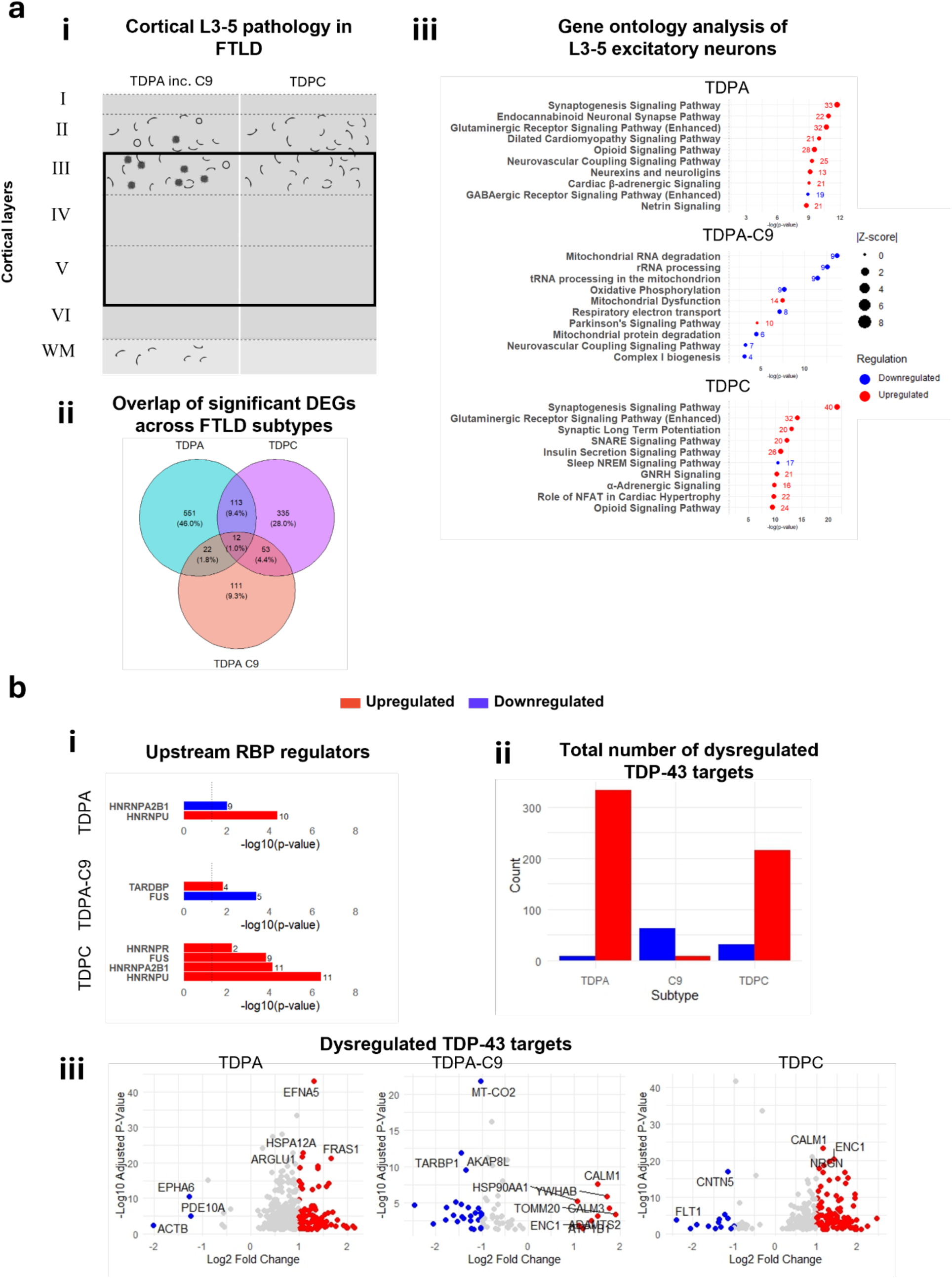
Characterisation of L3-5 excitatory neurons across FTLD-TDP subtypes: (a) (i) Schematic illustration of TDP-43 pathology in the cortex of FTLD-TDP subtype A (TDPA; n=3 and TDPA-C9; n=3) and subtype C (TDPC; n=4) cases, with a focus on L3-5 cortical layers. (ii) Venn diagram showing the overlap of significantly differentially expressed genes (DEGs) between FTLD-TDP subtypes relative to control in L3-5 excitatory neurons. Total gene counts are indicated, with the percentage of the total DEG pool shown in parentheses. (iii) Dot plot of the top 10 canonical pathways enriched in L3-5 excitatory neurons across FTLD-TDP subtypes compared to control. Pathway significance was determined by right-tailed Fisher’s exact test, with a -log₁₀ p-value threshold of 1.3 (dotted line). Dot size reflects the predicted activation z-score, while colour denotes activation status (red = positive, blue = negative). The number adjacent to each dot indicates the number of DEGs associated with that pathway. (b) (i) Upstream regulator analysis focused on heterogeneous nuclear ribonucleoproteins (hnRNPs) across FTLD-TDP subtypes. Regulators are ranked by -log₁₀ adjusted p-value, with direction of enrichment (up- or down-regulated) highlighted. NS = not significant. (ii) Bar plot showing the total number of DEGs overlapping with POSTAR3-derived TDP-43 targets, stratified by expression direction (red = upregulated, blue = downregulated). (iii) Volcano plots of differentially expressed TDP-43 target genes in each FTLD subtype (TDPA, TDPC, and TDPA with *C9orf72* mutation) relative to controls. Red and blue dots indicate significantly up- or downregulated genes, respectively, based on -log₁₀ adjusted p-value and log₂ fold change. Selected genes of interest are labelled.

These observations support the idea that pathological burden in superficial cortical layers drives more overt post-transcriptional alterations, whereas deeper-layer neurons may be more influenced by other RBPs and exhibit greater energy deficits and neurovascular changes, further highlighting TDP-43 pathology not being the only driving force of pathological changes across FTLD-TDP subtypes.

### Differential gene expression analysis reveals a predominant alteration of multiple hnRNPs in glial nuclei across FTLD subtypes

Further exploration of other hnRNP genes across various cell types was performed to gain deeper insight into alterations of the hnRNP network across FTLD subtypes (**Fig. 6a**). *TARDBP* expression was not significantly altered across any subtypes and cell clusters. In the TDP A vs CTRL comparison (**Fig. 6a i**), the most marked changes across cell clusters were primarily seen as transcriptomic increases of various HNRNP genes in mature oligodendrocytes, astrocytes, and mild changes occurring in endothelial cells, with a significant reduction in *HNRNPL* in astrocytes. With regards to TDP A neuronal populations, the only significant changes were seen in *HNRNPDL* in L2-3 excitatory neurons in the sporadic cases. For TDP A cases carrying C9-mutations (**Fig. 6a ii**), there were fewer clusters that attain significance, however, the majority of the changes seen in oligodendrocytes and the *HNRNPL* alterations seen in astrocytic cluster were still maintained. This was similarly reflected in other datasets (**Supp. Fig. 6**) where oligodendrocyte clusters exhibit widespread upregulation of HNRNP genes except for *HNRNPL*, and *HNRNPDL*, which was further downregulated across other clusters; astrocytes, OPCs, SST interneurons, and excitatory neurons (L2-3, L3-5, and L6). In the TDP C vs CTRL comparison (**Fig. 6a iii**), the hnRNP network was almost entirely upregulated in mature oligodendrocytes, with some other clusters; astrocytes, exhibiting upregulation of *HNRNPA2B1, HNRNPC, HNRNPDL, PCBP2*, and *RBMX* and OPCs exhibiting upregulation of *HNRNPA2B1* and *HNRNPU*. *HNRNPL* was downregulated in L2-3 neurons, mature oligodendrocytes, and OPCs, similar to patterns observed in TDP A and TDP A-C9.

**Fig. 6.**
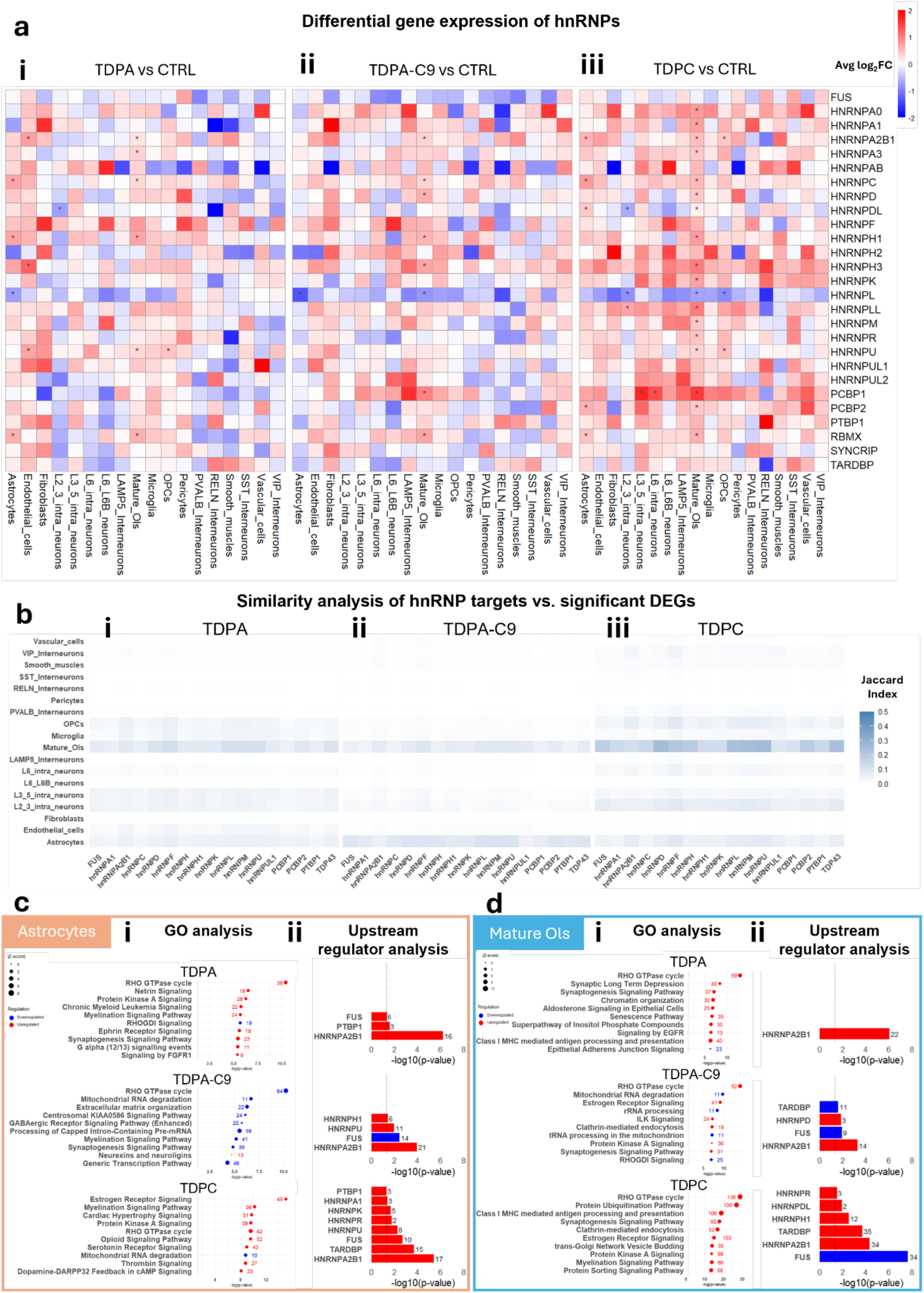
Transcriptional dysregulation of hnRNPs in glial clusters in FTLD subtypes: (a) Heatmaps showing the differential gene expression of FTLD (i, TDP A; n=3, ii, TDP A-C9; n=3, and iii, TDP C; n=4) cases against controls. Red- and blue-coloured boxes indicate a positive and negative log₂ fold change, respectively, with an asterisk indicating a false discovery rate (FDR) of <0.05. FDR was derived using Wilcox test followed by the Benjamini-Hochberg method (b) Similarity analysis heatmap of hnRNP targets with significant differential gene expression profile across all neural nuclei clusters of (i) TDP A, (ii) TDP A-C9, and (iii) TDP C. Higher Jaccard indices are indicated by more solid-coloured boxes. (c) (i) Dot plots illustrating the top 10 canonical pathways significantly enriched in astrocytes from FTLD-TDP cases relative to control. Enrichment was assessed using a right-tailed Fisher’s exact test, with a -log₁₀ p-value significance threshold of 1.3 (dotted line). Dot size corresponds to the activation z-score, and colour indicates activation state (red = activated; blue = inhibited). Numbers next to each dot represent the count of DEGs mapping to each pathway. (ii) Upstream regulator analysis constrained to hnRNP family members. Regulators are ranked by statistical significance (-log₁₀ adjusted p-value), and direction of enrichment is indicated. (d) (i) Dot plots illustrating the top 10 canonical pathways significantly enriched in mature oligodendrocytes from FTLD-TDP cases relative to control. Enrichment was assessed using a right-tailed Fisher’s exact test, with a -log₁₀ p-value significance threshold of 1.3 (dotted line). Dot size corresponds to the activation z-score, and colour indicates activation state (red = activated; blue = inhibited). Numbers next to each dot represent the count of DEGs mapping to each pathway. (ii) Upstream regulator analysis constrained to hnRNP family members. Regulators are ranked by statistical significance (-log₁₀ adjusted p-value), and direction of enrichment is indicated.

The disruption of the hnRNP network was further exemplified by exploring differential gene expression of the different targets of the hnRNPs and how similarly they overlap with each cluster (**Fig. 6b**). Similar to what was reported with the heatmaps, Jaccard similarity analyses revealed that there was an increased overlap between oligodendrocyte, astrocytes, and L2-3 excitatory neurons with most hnRNP targets (**Fig. 6b i and ii**). This was particularly pronounced in TDP C where astrocytes and OPCs also exhibited elevated Jaccard index scores (**Fig. 6b iii)**. Furthermore, TDP C oligodendrocyte cluster also showed higher Jaccard indices with HNRNPD, L, M, U, and TDP-43 targets.

These findings further underlined the important role non-neuronal cells have in FTLD and how that differs across the different subtypes. In astrocytes, pathway analysis highlights subtype-specific enrichments in signalling, structural, and metabolic processes, with TDP A showing strong synaptic and axon guidance pathways, TDP A-C9 exhibits changes in mitochondrial and transcriptional processes, and TDP C was enriched in neuro-modulatory signalling (**Fig. 6c i, Supp. Table 3)**. Further exploring the RBP network beyond transcriptional changes, upstream regulator analysis identified distinct sets of RBPs enriched in each subtype’s DEG profile. Across all three subtypes, *HNRNPA2B1* exhibited the most significant increase in z-scores. TDP C showed the broadest spectrum of changes, including increased *TARDBP, FUS, HNRNPU, R, K, A1,* and *PTBP1* (**Fig. 6c ii)**. In mature oligodendrocytes, pathway analysis reveals that TDP A is enriched in cytoskeletal, chromatin, and immune-related pathways, while C9 shows strong signals in RNA processing and mitochondrial dysfunction, and TDP C was characterised by changes in protein degradation, sorting, and antigen presentation (**Fig. 6d i)**. Similar to the neuronal and astrocytic clusters, upstream RBP regulator analysis shows that TDP C oligodendrocytes exhibit the broadest involvement of RBPs, including *TARDBP, FUS*, and several HNRNPs, suggesting deeper dysregulation of RNA metabolism in this subtype. TDP A oligodendrocytes with and without C9 mutations display a more restricted regulatory pattern, with *HNRNPA2B1* as the predominant factor (**Fig. 6d ii)**.

### Astrocytes and oligodendrocytes transcriptionally drive splice alterations in FTLD-TDP

To explore disease associated transcriptomic changes in the astrocytes and oligodendrocytes in more detail, and to explore the transcriptional heterogeneity underlying FTLD-TDP, we applied high-dimensional Weighted Gene Co-Expression Network Analysis (hdWGCNA). The analysis identified eight distinct eigengene co-expression modules, each characterised by unique transcriptional signatures within our dataset (**Fig. 7a**). These modules were named according to their assigned colours (e.g., turquoise, blue, pink) and were further analysed to understand their biological significance. Turquoise, black, and pink module eigengenes were enriched in neuronal clusters, yellow was highly expressed in microglia-assigned nuclei, and OPCs expressed red module eigengenes. Green and blue modules were expressed in astrocytes and mature oligodendrocytes, respectively, and the brown module was expressed in both.

**Fig. 7.**
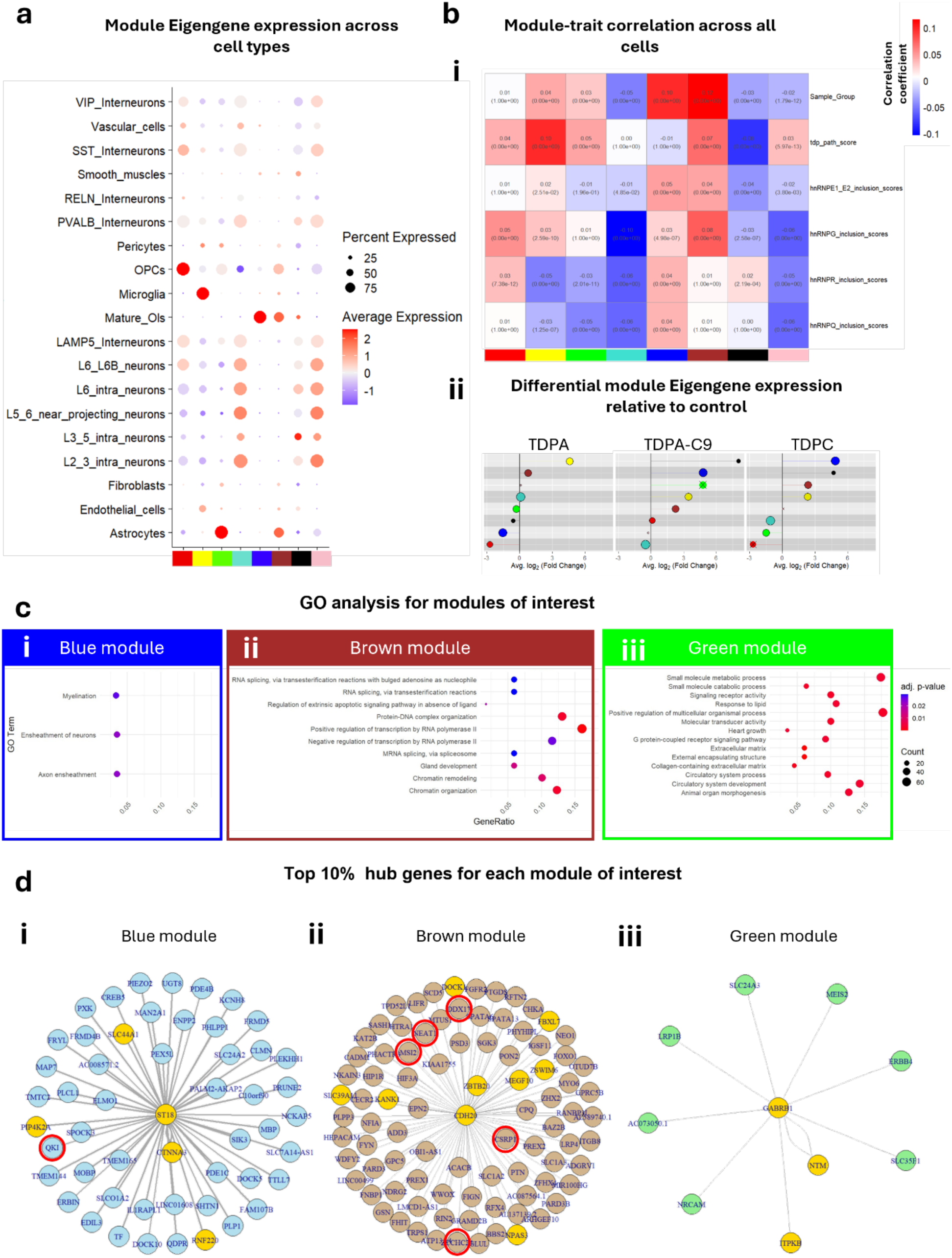
High-dimensional weighted gene co-expression network analysis of FTLD-TDP subtypes: (a) Dot plot summarising module eigengenes (MEs) identified through hdWGCNA, annotated by their enrichment across distinct cell types. (b) (i) Module–trait correlation matrix showing the relationship between module eigengene expression and clinical/pathological variables, including disease classification, TDP-43 pathology score, and hnRNP (E1/E2, G, R, Q) inclusion scores. Pearson’s correlation coefficients are visualised by colour intensity (red = positive, blue = negative); significance assessed using two-tailed *t*-tests with Bonferroni correction for multiple testing (FDR values shown in parentheses). (ii) Lollipop plots depicting changes in module eigengene expression for each FTLD-TDP subtype relative to control. Modules are plotted by average log₂ fold change; non-significant differences (FDR ≥ 0.05, Wilcoxon test with Benjamini–Hochberg correction) are marked with a cross. (c) Gene ontology (GO) enrichment analysis of selected modules: (i) blue, (ii) brown, and (iii) green. Dot plots display the top-ranked pathways by enrichment significance, with dot size indicating the number of genes contributing to each pathway. (d) Hub gene analysis for the same modules: (i) blue, (ii) brown, and (iii) green. The top 10% most interconnected genes within each module are shown, with the top 1% highlighted in yellow. Genes associated with RNA-binding, splicing, or transport functions are outlined in red.

To assess the relevance of gene co-expression modules to disease status and pathology, we first performed module-trait correlation analysis across all metacells (**Fig. 7b i**). This revealed that the green, brown, blue, and yellow modules were significantly correlated with disease classification (i.e. control vs FTLD), indicating that their eigengene expression systematically differs between FTLD and control samples, irrespective of subtype. In contrast, the turquoise, black, and pink modules exhibited negative correlations with disease classification. Beyond clinical grouping, the same analysis revealed distinct transcriptional associations with inclusion burden for TDP-43 and several hnRNP proteins (**Fig. 7b i**). The pink, brown, green, yellow, and red modules showed positive correlations with both TDP-43 pathology scores, with brown, blue, and yellow exhibiting further correlations with the inclusion scores for hnRNP E1/E2 and G, suggesting that these modules capture shared transcriptional programs responsive to RBP aggregation. The blue module also showed further correlations with hnRNP R, and Q inclusion scores. HnRNP G inclusion scores were negatively correlated with turquoise, black, and pink modules, enriched in neuronal clusters, potentially indicating repression or functional loss of this program in affected cells. Correlations with hnRNP E1/E2 and hnRNP Q were generally weaker across modules, possibly reflecting more diffuse or cell-type-specific effects not captured in this global analysis. Collectively, these findings suggest that a subset of modules respond to convergent RBP pathology, while others may reflect selective vulnerability to specific protein aggregates.

To better understand how individual FTLD subtypes contribute to these global transcriptional trends, we next examined differential module eigengene (DME) expression across TDP A, TDP A-C9, and TDP C relative to control (**Fig. 7b ii**). The brown and yellow modules were consistently upregulated across subtypes, while the blue and black modules were upregulated in TDP A-C9 and TDP C, but downregulated in TDP A, highlighting divergent transcriptional responses between genetically and sporadically driven forms of the disease. The green module presented a more nuanced profile as it was slightly downregulated in TDP A and TDP C but substantially upregulated (though not significantly) in TDP A-C9. This opposing direction of change resulted in a net positive correlation with disease status in the correlation analysis, resolving the apparent discrepancy between the two approaches. These findings underscore how subtype heterogeneity within FTLD can shape and sometimes obscure global disease-associated transcriptional signals, reinforcing the need to contextualise gene module dynamics within individual pathological profiles.

Considering the expression and enrichment of MEs, coupled with the glial transcriptional disruption of the RBP network, across FTLD subtypes, we performed gene ontology (GO) enrichment analysis to reveal different disease-related functional changes that were specific for modules highly associated with astrocytes and oligodendrocytes (**Fig. 7c**). GO analysis revealed that the blue module, which was mainly present in oligodendrocytes, was highly enriched in myelination pathways (**Fig. 7c i**). The brown module, which was present in oligodendrocytes and astrocytes, were associated with RNA splicing, transcription regulation, and chromatin organisation (**Fig. 7c ii**). The green module, which was enriched in just astrocytes, was highly associated with small molecule processing, altered receptor activity, and responses to lipids (**Fig. 7c iii**).

To explore what genes drove expression across each of these modules of interest, we performed hub gene analysis in the modules of interest (**Supp. Table 4**). We isolated the top 10% of hub genes based on their kME values and highlighted the top 1% across each module of interest. In the blue module, key hub genes included *ST18, CTNNA3, PIP4K2, SLC44A1, RNF220* (**Fig. 7d i**). Whilst no RBP genes or genes regulating RNA splicing were identified in the top 1% of hub genes in the network plot, we identified *QKI* in the top 10%. On the other hand, the brown module contained *CDH20, ZBTB20, MEGF10, ZSWIM6, NPAS3, KANK1, SLC39A1, DOCK1, FBXL7* in the top 1% hub genes (**Fig. 7d ii**). With a focus on RBP-related functions; *ZCCHC2, DDX17, CSRP1*, *MSI2,* and *NEAT1* were identified as important in relating pre-mRNA splicing, mRNA export from the nucleus, and mRNA stability. In the green module, *GABRB1, NTM, and ITPKB* were identified as the top hub genes, with no hub genes coding for a protein involved in RNA-binding (**Fig. 7d iii**).

## Discussion

Our data supports the role of multiple members of the hnRNP family in FTLD and further outlines the need to investigate non-neuronal cell types in these diseases. As seen with TDP-43, other hnRNPs show variable amounts of pathology in the FTLD frontal cortex and hnRNP protein accumulation was evident in multiple glial cell types. Changes in the hnRNP network in glial cell types were further confirmed at the transcriptomic level by snRNA-seq.

TDP-43 is the most extensively studied hnRNP in the context of FTLD, and its pathological aggregation forms the basis for subtyping FTLD-TDP cases. Subtype classification relies on immunohistochemical detection of pTDP-43 within distinct inclusions or neuritic structures, typically located in specific cortical layers of the frontal and temporal lobes. In this study, we demonstrate that the frequency of pTDP-43-positive inclusions and neurites in these cortical regions was highly variable, even among cases with the same pathological subtype. The hippocampal dentate gyrus (DG), a region also frequently affected in FTLD, exhibits NCIs and is impacted by pTDP-43 pathology. We confirm that in the DG, pTDP-43 pathology is restricted to NCIs, and we observe that type C cases, characteristically sporadic and associated clinically with SD (5), exhibit the highest frequency of NCIs within the granule cells. Conversely, sporadic type A cases, typically linked to bvFTD (5), show the lowest NCI burden in the DG. Notably, the frequency of pathological inclusions in the hippocampus does not correlate with the burden observed in the frontal and temporal cortices.

We then investigated immunohistochemical staining of other hnRNPs in FTLD frontal cortex cases from different sporadic (TDP A and TDP C) and genetic subtypes (TDP A-C9). We have previously observed the mislocalisation of hnRNP R and Q into pathological inclusions in FTLD-FUS patient tissue (46,47) and we herein observe their accumulation in axonal and neuritic projections in FTLD-TDP subtypes as well. We also confirm the presence of hnRNP E1/2 in dystrophic neurites typical of TDP C (21,22). Whilst multiple hnRNPs may not form *bona fide* pathological inclusions, we observed a shift in the localisation of multiple hnRNP members, with a greater predominance in the cytoplasm of neurons in FTLD, an alteration that could itself contribute to aberrant protein function. This further supports the concept of hnRNPs functioning as a coordinated network, where the aggregation of one member may disrupt the distribution or function of others (25).

We observed an increase in hnRNP immunoreactivity in non-neuronal cells. An increase in hnRNP G was observed in astrocytes across all FTLD subtypes. HnRNP G has been identified as having a central role in the DNA damage response pathway and was reported to be acutely upregulated in astrocytes post-injury in preclinical spinal-cord injury models (48–50). HnRNP R and Q showed subtype-specific aggregation, with hnRNP Q also being observed accumulated in astrocytes (TDP A and C) and microglia (TDP C). Importantly, hnRNP Q and R are known to modulate TDP-43 toxicity and RNA binding, with *SYNCRIP* knockdown shown to rescue TDP-43-induced toxicity in model systems (51). HnRNP Q is also dysregulated in ALS, displaying distinct subcellular localisation in sporadic vs. *C9orf72* cases (52). We have shown that both hnRNP Q and hnRNP R were found in pathological inclusions in FTLD-FUS (46), although their functional relationship remains unclear. Our observed glial redistribution of hnRNP R and Q may therefore reflect broader molecular disturbances. This aligns with upstream regulator analyses predicting dysregulated FUS activity in glial populations across FTLD-TDP subtypes. Together, these findings support the hypothesis that glial hnRNP dysregulation may contribute to disease pathogenesis through networked interactions with pathological proteins such as TDP-43 and FUS (53).

SnRNA-seq analysis on the same cases revealed that changes within the hnRNP protein network can be fairly different to what emerges at the transcriptomic level. Our analysis identified *HNRNPL* as the only hnRNP whose expression was significantly altered in at least one cluster across all subtypes, which was confirmed in the Gittings dataset (**Supp. Fig. 5**; (31). Previous reports have identified hnRNP L as a stabilising factor that protects vulnerable transcripts from degradation by nonsense-mediated decay (NMD), particularly those with long 3′ UTRs (54,55). In parallel, hnRNP L also functions as a splicing regulator of *UNC13A* (*56*), a key downstream target of TDP-43, and its nuclear levels have been shown to increase *in vitro* in response to TDP-43 aggregation (57). Despite the observed alterations in *HNRNPL* gene expression, we did not identify significant abnormal localisation or immunohistochemical patterns of the HnRNP L protein in FTLD. Conversely, *TARDBP* itself did not exhibit significant differential expression in any cell cluster, but does show protein inclusions and mislocalisation, further emphasising the complexity of hnRNP involvement in disease at the transcriptomic and protein level.

Upstream analysis of the snRNA-seq data predicted reduced FUS protein activity in L2–3 neurons across FTLD-TDP subtypes, despite no direct evidence of FUS aggregation or mislocalisation, suggesting that these changes may reflect functional impairments secondary to TDP-43 pathology or a loss of FUS function that occurs without detectable changes in protein abundance. These findings hint at subtype-specific mechanisms of FUS dysregulation, possibly stemming from the distinct structural and biochemical properties of TDP-43 aggregates. Type A inclusions, for instance, are more insoluble and sequester more RNA-binding proteins than the structurally less stable type C neurites, which have milder biochemical effects (58–61). Cryo-EM studies have shown that type A fibrils resist degradation and disrupt RNA-protein interactions, impairing nucleocytoplasmic transport via interference with transportin-1 (TNPO1), a key FUS importer (62). This may explain FUS-associated gene dysregulation. Notably, analysis of hnRNP targets revealed greater transcriptional overlap in TDP C, implying broader but less severe RNA network disruption, whereas TDP A cytoplasmic inclusions appear to drive more localized dysfunction through greater hnRNP sequestration (58–61).

Furthermore, our analysis of pathway-level alterations in L2-3 excitatory neurons further supports the view that post-transcriptional vulnerabilities are compounded by failures in translational control and stress response signalling, particularly via suppression of EIF2AK4 (GCN2)-mediated integrated stress response. This suppression, most prominent in sporadic TDP A and C, was accompanied by reduced activity across core protein synthesis pathways, including translation initiation, elongation, termination, and nonsense-mediated decay. Given EIF2AK4’s role in modulating translation under nutrient and ribosomal stress (63), these disruptions suggest a failure to maintain proteostatic balance. Although EIF2AK4 dysregulation has been reported in *C9orf72* models (64), our findings in sporadic cases highlight its broader relevance and the importance of human tissue studies. Notably, this translational repression was largely confined to L2-3 neurons, consistent with their heightened vulnerability to TDP-43 pathology. In contrast, L3-5 neurons displayed more variable subtype-specific changes. While deep-layer neurons showed fewer consistent DEGs, they nonetheless revealed distinct subtype patterns. Notably, in TDP A, L3-5 neurons exhibited upregulation of synaptic and neurovascular signalling, including glutamatergic, GABAergic, endocannabinoid, and neurexin/neuroligin pathways, suggestive of a shift toward synaptic remodelling and altered excitatory-inhibitory balance (65). Additionally, increased ROBO signalling in TDP A and C may reflect compensatory or maladaptive responses to axonal injury, as previously implicated in ALS (66–68).

Together, our findings further underscore a layer-specific pattern of TDP-43–associated dysfunction, with L2-3 neurons exhibiting weaker enrichment of other RBPs with greater disruption of TDP-43 target transcripts in TDP C. In contrast, L3-5 neurons show a higher signature of RNA dysregulation, with subtype-specific differences in both upstream RBPs and elevated downstream TDP-43-related transcript profiles. These observations support the hypothesis that pathological burden in superficial cortical layers drives more overt post-transcriptional alterations, whereas deeper-layer neurons may be more related to other RBPs and exhibit greater energy deficits and neurovascular changes.

Beyond neuronal populations, our snRNA-seq analysis highlights glial disruptions in the hnRNP network. TDP C cases show pronounced gene expression changes in oligodendrocyte clusters compared to other subtypes. To support these observations, we reanalysed hnRNP expression using two independent published datasets (31,69). Indeed, similar to our own findings, previous snRNA-seq studies performed on FTLD-GRN reported significant astrocytic changes, including transcriptomic disruptions in genes linked to increased synaptic degeneration (70). Together, these findings highlight the contribution of glial cells to FTLD-TDP pathophysiology and suggest hnRNP alterations may converge with broader astrocytic dysfunction in disease.

Similar to neuronal populations, glial populations also showed consistent hnRNP dysregulation via the upstream regulator analysis. The most consistent disruption was in hnRNP A2B1 across all FTLD-TDP subtypes in astrocytes and oligodendrocytes. HnRNP A2B1, a regulator of RNA splicing and transport essential for oligodendrocyte and astrocyte function, can become dysregulated in response to neuronal TDP-43 pathology, as these neurons fail to maintain proper neuron-glia communication (19,71). This was further reflected in the co-expression network analysis, where glial-enriched modules were found to associate robustly with disease classification, aligning with recent reports (72). Notably, NEAT1, an lncRNA involved in paraspeckle formation and stress-responsive splicing regulation (73), emerges as a key player, along with RBPs such as MSI2, DDX17, and QKI. QKI, in particular, functions as a hub in oligodendrocyte-enriched modules, linking RNA processing deficits to disrupted myelination and glial maturation (74–77). While the astrocyte module lacked classical RNA-processing hubs, it exhibited signatures of impaired small molecule and lipid metabolism. These findings support a model in which glial cells contribute to FTLD-TDP pathogenesis through cell-type-specific reprogramming of RNA regulation, potentially reflecting both compensatory and maladaptive responses to upstream RBP dysfunction.

Increasing age is also known to cause alterations in hnRNP expression in a cell-specific manner, which has consequences on downstream RNA processing, resulting in more age-related mis-splicing of downstream targets (78). Moreover, a recent study has found that ageing can be sufficient to trigger mislocalisation of RBPs such as TDP-43 *in vitro*, potentially through the induction of a chronic cellular stress cycle which leads to a reduction of RBP nuclear presence (79). Our study compares age-matched controls to FTLD cases and therefore highlights that there are glial-specific gene expression changes within the hnRNP network that may be specific or exacerbated in disease. However, these changes might not lead to changes in the protein location or the formation of pathological inclusions. This disconnect between pathological markers and transcriptomic alterations has been observed in a recent study, where cryptic exon expression correlated more strongly with each other than with TDP-43 pathology in the hippocampus (80). Overall, whilst some transcriptomic changes likely reflect downstream effects of pathology (e.g. mitochondrial stress; (31,81), others linked to RNA and synaptic signalling genes might be a direct consequence of TDP-43 mislocalisation, especially in L2-3 neurons.

In conclusion, our study contributes towards FTLD pathological subtype-specific transcriptomic changes using snRNA-seq and the characterisation of pathological changes within the HnRNP network via histological studies. Importantly, we characterise changes in hnRNP expression at both the transcriptomic and protein level within non-neuronal cells. These changes were particularly pronounced in the oligodendrocyte lineage in our snRNA-seq data. Our findings suggest non-neuronal cell types and the pathological characterisation of other HnRNPs should be taken into consideration when investigating pathological and transcriptomic changes occurring in FTLD.

## Acknowledgements

We would like to thank Carlo Sala Frigerio (previously University College London) for his contribution to study design and the execution of the single-nuclei RNA sequencing experiment. The Queen Square Brain Bank is supported by the Reta Lila Weston Institute of Neurological Studies, UCL Queen Square Institute of Neurology.

## Funding Declaration

TL and AG are supported by Alzheimer’s Society. TL is supported by Alzheimer’s Research UK and the Association of Frontotemporal Dementia. AG is supported by BRAIN non-clinical fellowship and My Name’5 Doddie Foundation. YB is supported by the Association of Frontotemporal Dementia.

## Ethics Approval

The procurement and use of human tissues in this study was in accordance with the UK Human Tissue Act 2004. All samples were supplied, anonymised by Queen Square Brain Bank, UCL Queen Square Institute of Neurology and had full research consent (REC 23/LO/0044).

## Author contributions

A.G, Y.B, and T.L conceptualised and designed the experiments. Y.B. analysed snRNA-seq data and visualised results. J.H. validated snRNA-seq sample IDs. T.L., A.G., B.B., L.G., P.G-P., S.F and B.F. performed the immunohistochemical analysis. T.L. and A.G. generated neuropathological data. T.L., A.G., Y.B., K.F., and C.T. interpreted the data. All authors read and approved the final version of the manuscript.

## Data availability

The raw and pre-processed snRNA-seq data in this publication have been deposited in NCBI’s Gene Expression Omnibus and are accessible through GEO series accession number GSE288106.

## Competing interest

The authors declare no conflicts of interest

## Supplementary tables and figures

**Supplementary Table 1:** Full demographic data of all the cases used in the study, including a heatmap of variable phospho-TDP-43 pathology. AAO = age at onset. AAD = age at death. In the pathology heatmap, blue cells indicate minimal pathology whilst red cell indicate high levels of pathology.

**Supplementary Table 2:** Summary table of the number of nuclei collected for single-nuclei RNA sequencing across each case and how they distribute across different clusters with differential gene expression results across all cell types

**Supplementary Table 3:** Summary table of the canonical pathways enriched across cell types in the single-nuclei RNA sequencing dataset.

**Supplementary Table 4:** Module eigengene based connectivity across modules in the FTLD-TDP snRNAseq dataset

**Supplementary Figure 1:** Demographics comparison against TDP-43 pathological scoring in the (a) temporal cortex, (b) frontal cortex, and (c) hippocampal granule cell layer. Botplots contain the comparison of FTLD subtypes for (d) age of onset in years, (e) age at death in years, (f) disease duration in years, and (g) post-mortem interval in hours. Data were analysed using a simple linear regression analysis and Kruskal Wallis analysis, respectively. ** indicates p<0.01.

**Supplementary Figure 2:** HnRNP immunohistochemistry and quantification in different FTLD subtypes. Immunohistochemical images of HnRNP A1, A2B1, C1/2, D and E1/2 staining in the frontal cortex in all FTLD subtypes as well as controls. The violin plots next to the corresponding HnRNP images depict the values listed in Table 3. Scale bar indicates 50µm. In all plots N=Nuclear, C= Cytoplasmic, I= inclusions.

**Supplementary Figure 3:** HnRNP immunohistochemistry and quantification in different FTLD subtypes. Immunohistochemical images of HnRNP F, G, H, I, and L staining in the frontal cortex in all FTLD subtypes as well as controls. The violin plots next to the corresponding HnRNP images depict the values listed in Table 3. Scale bar indicates 50µm. In all plots N=Nuclear, C= Cytoplasmic, I= inclusions.

**Supplementary Figure 4:** HnRNP immunohistochemistry and quantification in different FTLD subtypes. Immunohistochemical images of HnRNP M, P, Q, R and U staining in the frontal cortex in all FTLD subtypes as well as controls. The violin plots next to the corresponding HnRNP images depict the values listed in Table 3. Scale bar indicates 50µm. In all plots N=Nuclear, C= Cytoplasmic, I= inclusions.

**Supplementary Figure 5:** Most significant DEGs in different cell types across different FTLD subtypes: Heatmaps displaying log₂ fold change of the top 75 most significantly differentially expressed genes in L2-3 and L3-5 excitatory neurons, across FTLD subtypes (TDP A, TDP A-C9, and TDP C) compared to controls.

**Supplementary Figure 6:** SnRNA-seq analysis of the genes belonging to the HnRNP network from the published datasets: Heatmaps showing the differential gene expression of TDP A-C9 cases against controls from Gittings et al. (2023) and Li et al. (2023). Red- and blue-coloured boxes indicate a positive and negative log₂ fold change, respectively, with an asterisk indicating a false discovery rate of <0.05.

## Notes

### Competing Interest Statement

The authors have declared no competing interest.

