## Supplementary figures and images for "Transcriptomic and pathological analysis of the hnRNP network reveals glial involvement in FTLD pathological subtypes"

### Supp fig 1

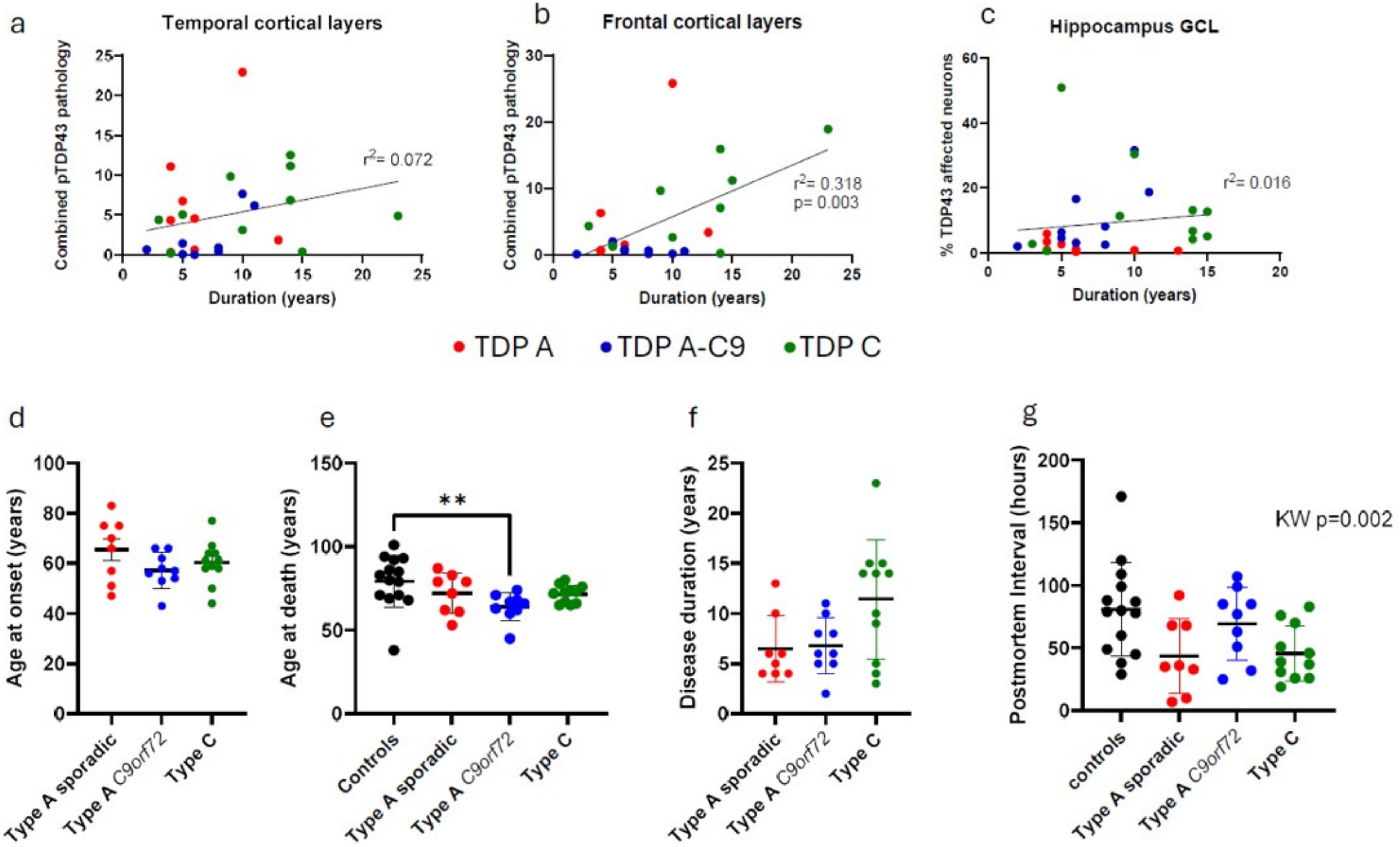

### Supp fig 2

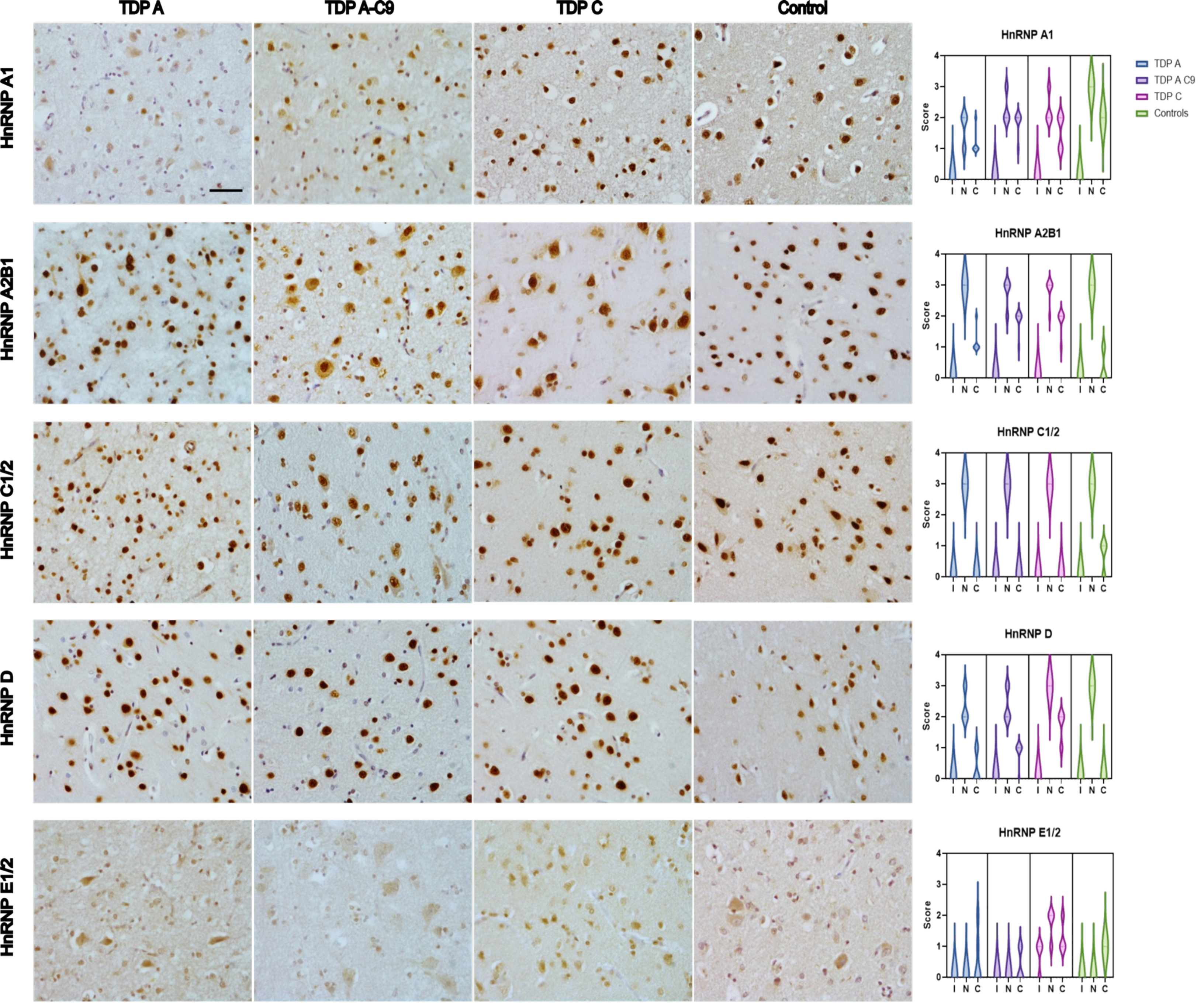

### Supp fig 3

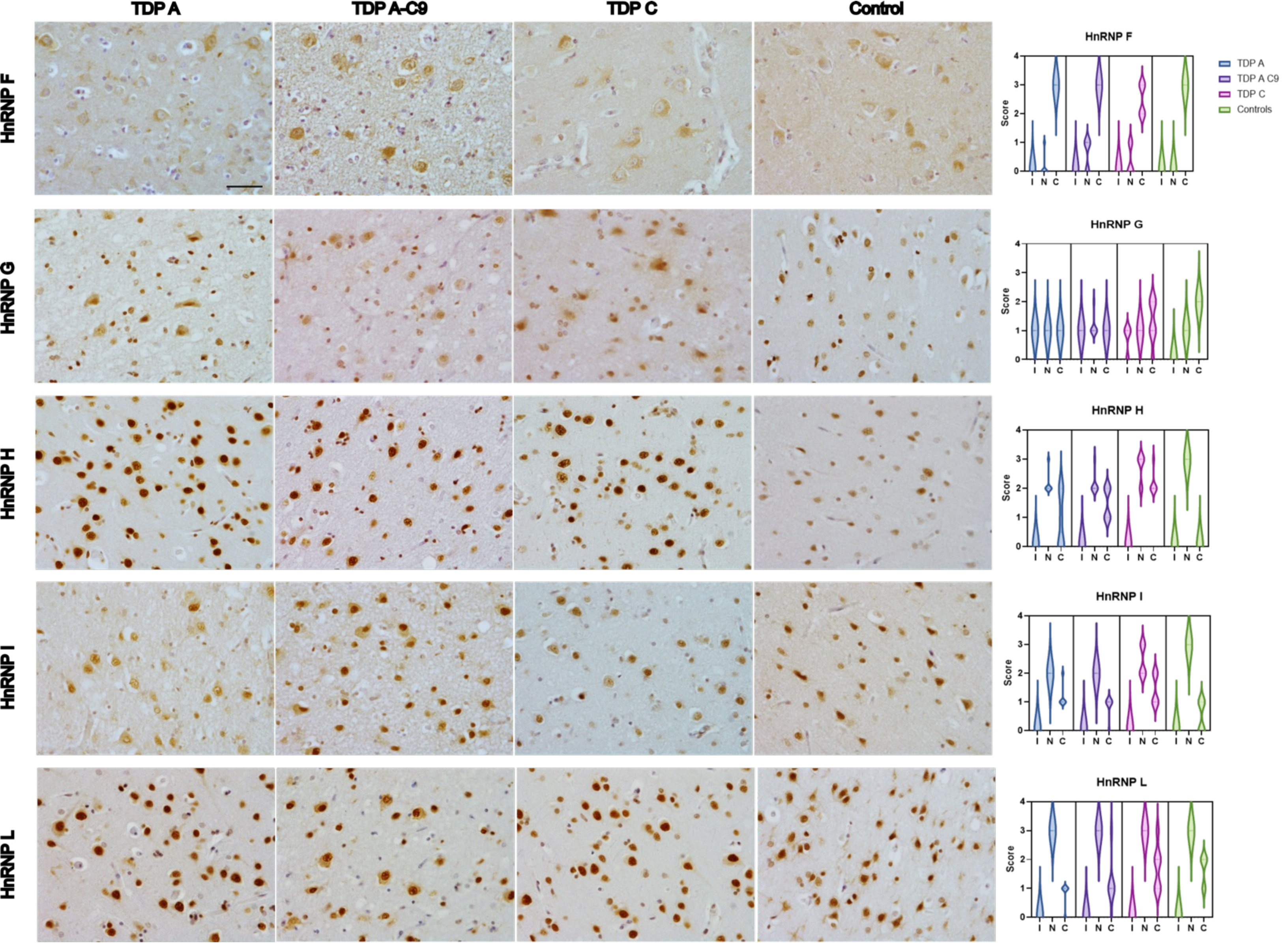

### Supp fig 4

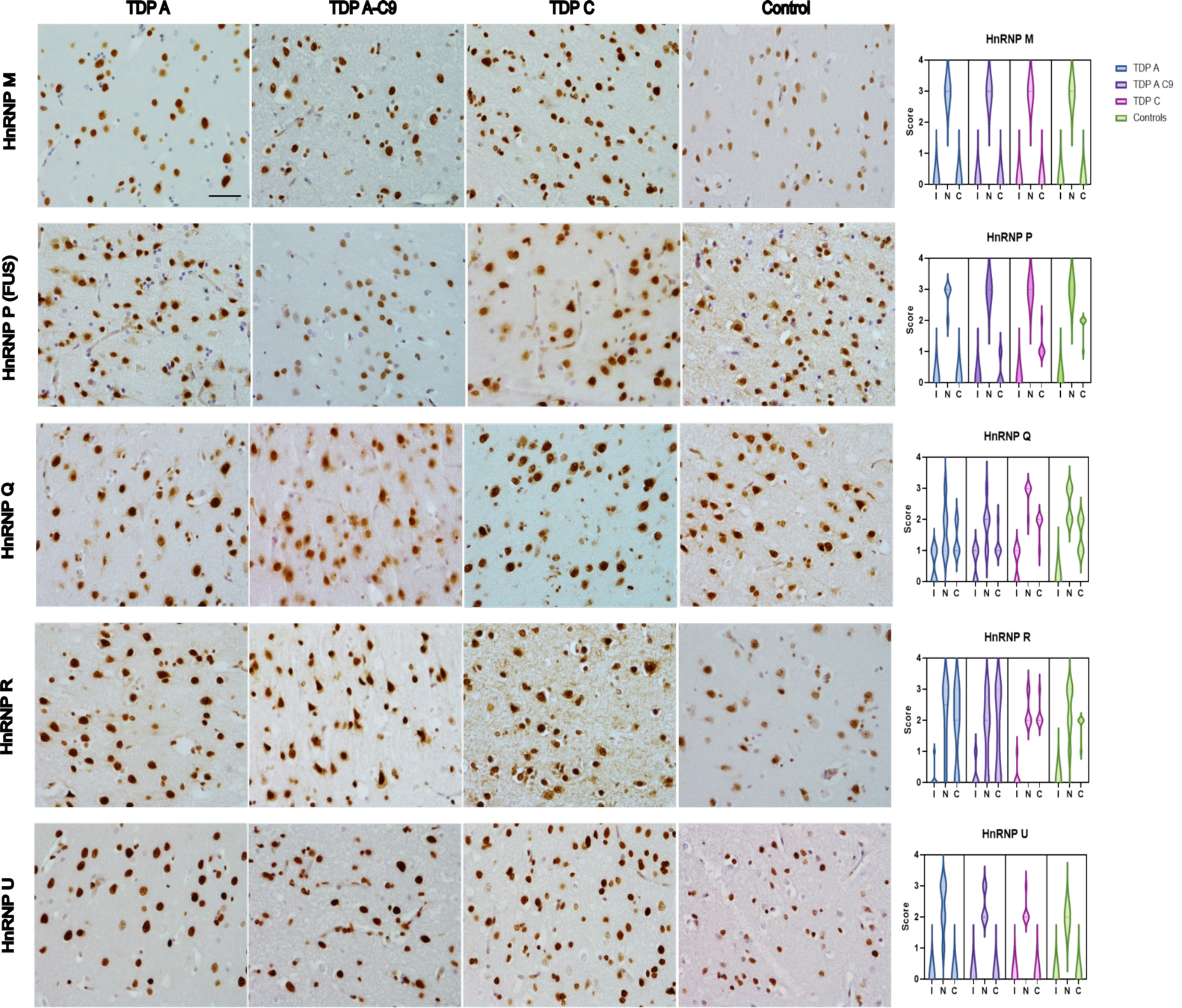

### Supp fig 5

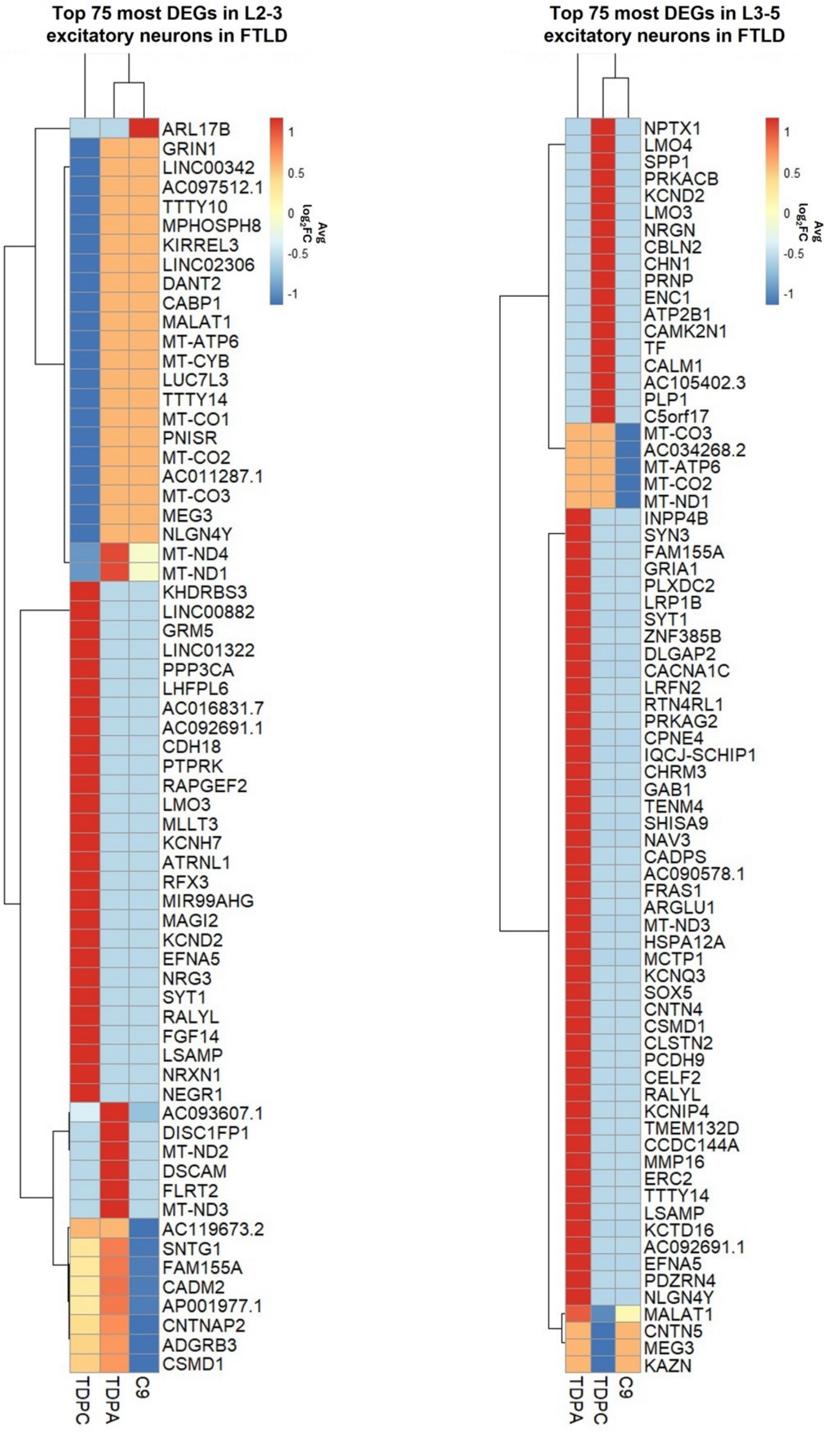

### Supp fig 6

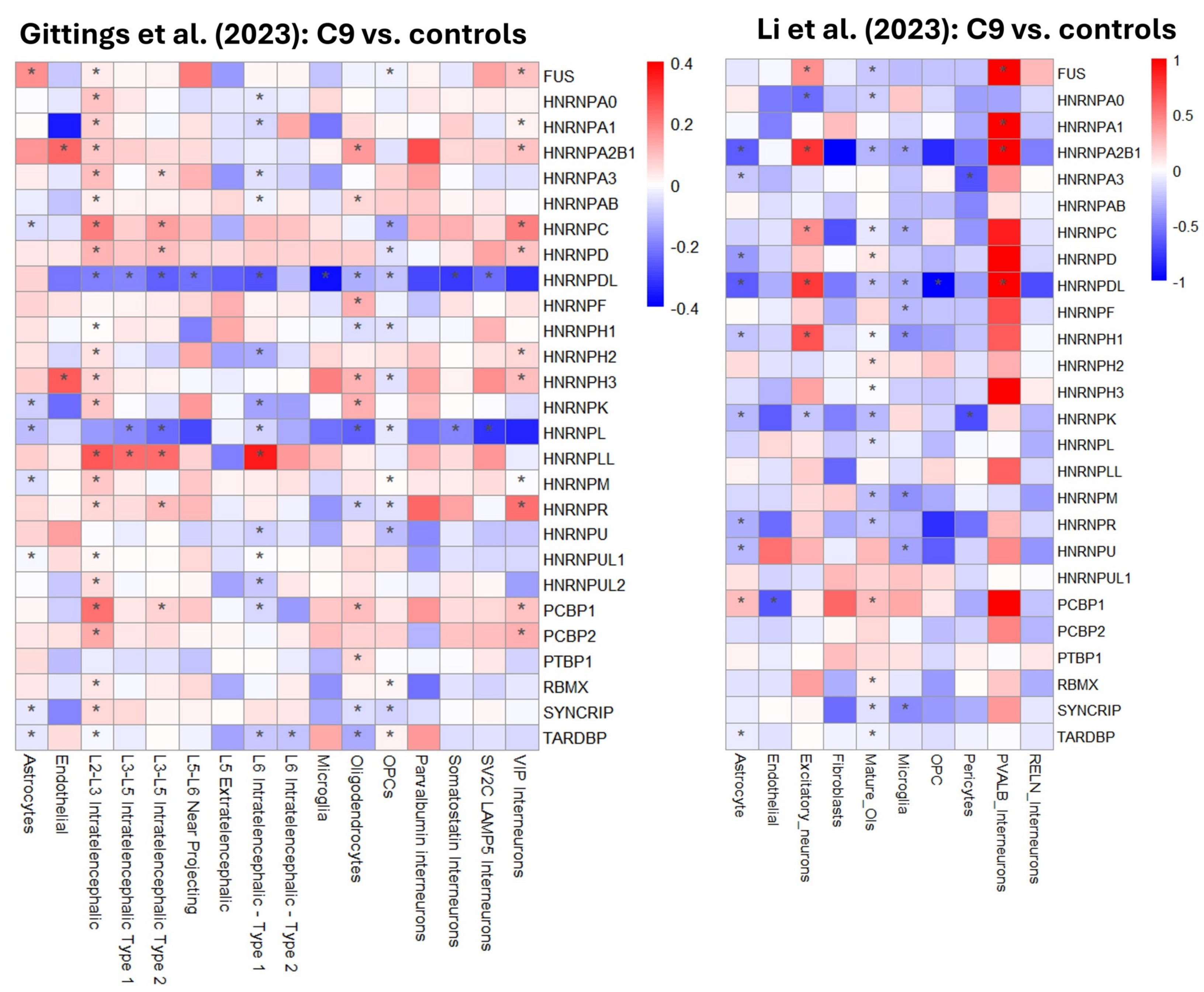
