## Supplementary material for "Transcriptomic and pathological analysis of the hnRNP network reveals glial involvement in FTLD pathological subtypes": Supp Table1

### Supp.Table 1

| Case no | Path Diag | Mutations | AAO (y) | AAD (y) | Disease duration (y) | Sex | % TDP43 affected neurons Hippocampus | Frontal cortex inclusions | Frontal cortex neurites | Frontal cortex pathology (combined) | Temporal cortex inclusions | Temporal cortex neurites | Temporal cortex pathology (combined) | Included in HnRNP immunohistochemical study | Included in ScRNASeq analysis |
|---|---|---|---|---|---|---|---|---|---|---|---|---|---|---|---|
| 1 | TDP A | | 57 | 62 | 5 | M | 2.6 | 0.8 | 1.1 | 1.9 | 1.4 | 5.3 | 6.7 | ✓ | |
| 2 | TDP A | | 51 | 61 | 10 | M | 0.7 | 1.8 | 24.0 | 25.8 | 2.7 | 20.2 | 22.9 | | ✓ |
| 3 | TDP A | | 66 | 72 | 6 | M | 0.2 | 0.8 | 0.8 | 1.5 | 0.5 | 4.1 | 4.6 | | ✓ |
| 4 | TDP A | | 77 | 79 | 2 | M | 0.9 | 0.3 | 0.4 | 0.7 | 0.3 | 10.8 | 11.1 | ✓ | |
| 5 | TDP A | | 70 | 83 | 13 | F | 0.6 | 2.9 | 0.4 | 3.4 | 1.6 | 0.3 | 1.9 | ✓ | |
| 6 | TDP A | | 75 | 79 | 4 | F | 5.8 | 0.8 | 5.5 | 6.3 | 0.1 | 0.2 | 0.3 | ✓ | |
| 7 | TDP A | | 47 | 53 | 6 | M | 1.1 | 0.1 | 0.6 | 0.7 | 0.6 | 0.0 | 0.6 | ✓ | ✓ |
| 8 | TDP A | | 83 | 87 | 4 | F | 3.4 | 0.1 | 0.5 | 0.6 | 1.8 | 2.5 | 4.4 | ✓ | |
| 18 | TDP A | *C9orf72* | 56 | 67 | 11 | F | 18.6 | 0.4 | 0.1 | 0.5 | 1.6 | 4.6 | 6.2 | ✓ | |
| 19 | TDP A | *C9orf72* | 54 | 60 | 6 | M | 3.0 | 0.7 | 0.1 | 0.8 | | | | ✓ | |
| 20 | TDP A | *C9orf72* | 57 | 62 | 5 | F | 6.3 | 1.1 | 0.4 | 1.5 | 1.3 | 0.2 | 1.4 | ✓ | ✓ |
| 21 | TDP A | *C9orf72* | 53 | 63 | 10 | M | 31.6 | 0.2 | 0.0 | 0.2 | 2.0 | 5.6 | 7.6 | ✓ | ✓ |
| 22 | TDP A | *C9orf72* | 66 | 74 | 8 | F | 2.4 | 0.0 | 0.6 | 0.6 | 0.2 | 0.2 | 0.4 | ✓ | |
| 23 | TDP A | *C9orf72* | 62 | 68 | 6 | M | 16.5 | 0.0 | 0.6 | 0.6 | 0.0 | 0.0 | 0.0 | ✓ | |
| 24 | TDP A | *C9orf72* | 43 | 45 | 2 | M | 1.9 | 0.1 | 0.0 | 0.1 | 0.6 | 0.0 | 0.6 | ✓ | |
| 25 | TDP A | *C9orf72* | 66 | 71 | 5 | M | 4.5 | 1.6 | 0.4 | 2.0 | 0.1 | 0.0 | 0.1 | | ✓ |
| 26 | TDP A | *C9orf72* | 58 | 66 | 8 | F | 8.0 | 0.2 | 0.0 | 0.2 | 0.9 | 0.0 | 0.9 | | |
| 27 | TDP C | | 77 | 80 | 3 | F | 2.7 | 0.9 | 3.4 | 4.3 | 1.8 | 2.6 | 4.4 | | |
| 28 | TDP C | | 64 | 78 | 14 | M | 6.6 | 0.0 | 0.2 | 0.2 | 6.9 | 4.3 | 11.1 | ✓ | ✓ |
| 29 | TDP C | | 58 | 72 | 14 | F | 13.1 | 0.0 | 15.9 | 15.9 | 0.0 | 12.5 | 12.5 | | ✓ |
| 30 | TDP C | | 66 | 76 | 10 | M | 11.3 | 0.0 | 9.6 | 9.7 | 0.1 | 9.7 | 9.8 | | ✓ |
| 31 | TDP C | | 50 | 65 | 15 | M | 12.6 | | | | | | | ✓ | |
| 32 | TDP C | | 61 | 66 | 5 | M | 50.9 | 0.0 | 1.3 | 1.3 | 0.2 | 4.8 | 5.1 | ✓ | |
| 33 | TDP C | | 58 | 73 | 15 | F | 5.0 | 0.2 | 11.0 | 11.2 | 0.0 | 0.4 | 0.4 | ✓ | ✓ |
| 34 | TDP C | | 64 | 74 | 10 | M | 30.3 | 0.0 | 2.6 | 2.6 | 0.0 | 3.1 | 3.1 | ✓ | ✓ |
| 35 | TDP C | | 59 | 73 | 14 | F | 4.1 | 0.1 | 7.0 | 7.1 | 0.7 | 6.2 | 6.8 | ✓ | |
| 36 | TDP C | | 44 | 67 | 23 | M | 46.8 | 0.4 | 18.5 | 18.9 | 0.5 | 4.4 | 4.9 | ✓ | |
| 37 | TDP C | | 60 | 65 | 5 | M | 0.5 | | | | 0.1 | 0.1 | 0.2 | ✓ | |
| 38 | Control | | n/a | 79 | n/a | F | 0 | 0 | 0 | 0 | 0 | 0 | 0 | ✓ | |
| 39 | Control | | n/a | 69 | n/a | M | 0 | 0 | 0 | 0 | 0 | 0 | 0 | ✓ | ✓ |
| 40 | Control | | n/a | 38 | n/a | M | 0 | 0 | 0 | 0 | 0 | 0 | 0 | ✓ | |
| 41 | Control | | n/a | 94 | n/a | F | 0 | 0 | 0 | 0 | 0 | 0 | 0 | ✓ | |
| 42 | Control | | n/a | 85 | n/a | M | 0 | 0 | 0 | 0 | 0 | 0 | 0 | ✓ | |
| 43 | Control | | n/a | 92 | n/a | F | 0 | 0 | 0 | 0 | 0 | 0 | 0 | ✓ | |
| 44 | Control | | n/a | 71 | n/a | F | 0 | 0 | 0 | 0 | 0 | 0 | 0 | | ✓ |
| 45 | Control | | n/a | 86 | n/a | F | 0 | 0 | 0 | 0 | 0 | 0 | 0 | | |
| 46 | Control | | n/a | 68 | n/a | F | 0 | 0 | 0 | 0 | 0 | 0 | 0 | | |
| 47 | Control | | n/a | 80 | n/a | F | 0 | 0 | 0 | 0 | 0 | 0 | 0 | | ✓ |
| 48 | Control | | n/a | 93 | n/a | F | 0 | 0 | 0 | 0 | 0 | 0 | 0 | | |
| 49 | Control | | n/a | 83 | n/a | F | 0 | 0 | 0 | 0 | 0 | 0 | 0 | | ✓ |
| 50 | Control | | n/a | 71 | n/a | M | 0 | 0 | 0 | 0 | 0 | 0 | 0 | | |
| 51 | Control | | n/a | 101 | n/a | M | 0 | 0 | 0 | 0 | 0 | 0 | 0 | | ✓ |

Key: 0 10 20 30 40 50 60
